# Characterization of essential eggshell proteins from *Aedes aegypti* mosquitoes

**DOI:** 10.1101/2020.04.06.027706

**Authors:** Carter J. Simington, Max E. Oscherwitz, Alyssa J. Peterson, Alberto A. Rascón, Brooke B. Massani, Roger L. Miesfeld, Jun Isoe

**Affiliations:** Department of Chemistry and Biochemistry, The University of Arizona, Tucson, AZ 85721; Department of Chemistry, San José State University, San José, CA 95192

**Keywords:** mosquito, eggshell, RNA interference, melanization, reproduction

## Abstract

Up to 40% of the world population now live in areas where dengue mosquito vectors coexist with humans. *Aedes aegypti* are vectors for zoonotic diseases that affect hundreds of millions of individuals per year globally. We recently identified the eggshell organizing factor 1 (EOF1) protein using systematic RNA interference (RNAi) screening of mosquito lineage-specific genes. It was shown that eggs deposited by RNAi-EOF1 *A. aegypti* and *A. albopictus* mosquitoes were non-melanized, fragile, and contained nonviable embryos. Motivated by this discovery, we performed RNAi screening of eggshell proteins to determine putative downstream target proteins of intracellular EOF1. We identified several eggshell proteins as essential for eggshell formation in *A. aegypti* and characterized their phenotypes in detail by molecular and biochemical approaches. We found that Nasrat, Closca, and Polehole structural proteins, together with the Nudel serine protease, are indispensable for eggshell melanization and egg viability. While all four proteins are predominantly expressed in ovaries of adult females, Nudel mRNA expression is highly upregulated in response to blood feeding. Furthermore, we identified four secreted eggshell enzymes as important factors for controlling the processes of mosquito eggshell formation and melanization. These enzymes included three dopachrome converting enzymes and one cysteine protease. All eight characterized eggshell proteins were required for intact eggshell formation. However, their surface topologies in response to RNAi did not phenocopy the effect of RNAi-EOF1. Still, it remains unclear how EOF1 influences eggshell formation and melanization. The use of proteomic analysis of eggshell proteins from RNAi-EOF1 assisted in the identification of additional proteins that could be regulated in EOF1 deficient eggshells.

## Introduction

Mosquito-borne diseases such as dengue, Zika, yellow fever, and chikungunya are transmitted to humans via *Aedes aegypti* mosquitoes; we coexist with this mosquito species in many inhabitable areas of the world. There is an ongoing need to develop unique small molecule inhibitors, as mosquito insecticide resistance has become a serious problem worldwide (Murray et al., 2013; Bhatt et al., 2013; Fauci and Morens, 2016; Carrasco et al., 2019). It is important that control agents should be developed to specifically target mosquito-selective biochemical processes in order to decrease potential negative off-target effects on other organisms. In particular, humans and insects that are beneficial should not be inadvertently harmed by this strategy.

Once *A. aegypti* female mosquitoes ingest blood, they initiate early oogenesis. Mosquitoes contain ∼100 ovarioles per ovary, which are composed of primary and secondary follicles along with a germarium. Mosquito ovarian follicles develop synchronously throughout oogenesis (Hagedorn et al., 1977; Lea et al., 1978; Clements and Boocock, 1984; Raikhel et al., 2002; Uchida et al., 2004; Isoe and Hagedorn, 2007; Clifton and Noriega, 2011). The spherical follicle gradually transforms into an ellipsoid form by accumulating vitellogenin yolk proteins. A single layer of follicular epithelial cells surrounding the oocyte is responsible for secreting a majority of eggshell structural components between 18- and 54-hours post blood meal at 28°C. Thus, the timing of biosynthesis, secretion, and formation of eggshell components during oocyte maturation is crucial for completing embryonic development after oviposition.

In general, the eggs of insects are sensitive to desiccation. Due to environmental and other external factors, newly deposited wild type mosquito eggs are no exception to this trend/ pattern, which may impact the reproductive success of a species. Thus to combat this issue, when mosquito eggs are laid, the vitelline envelope protein-containing endochorion layer rapidly undergoes maturation by tanning and hardening processes catalyzed by several enzymes. The formation of an embryo-derived extracellular serosal cuticle that protects developing embryo and larva from the dangers associated with desiccation (Farnesi et al., 2017; Rezende et al., 2008; Vargas et al., 2014). *A. aegypti* eggshell proteins were discovered more than 25 years ago (Lin et al., 1993; Edwards et al., 1998), and several key eggshell enzymes involved in eggshell melanization and protein cross-linking have been characterized (Li, 1994; Ferdig et al., 1996; Han et al., 2000; Johnson et al., 2013; Fang et al., 2002; Kim et al., 2005; Li and Li 2005; Li and Li 2006). Moreover, proteomic studies have been performed on mosquito eggshells to identify the most abundant protein components (Amenya et al., 2010; Marinotti et al., 2014). However, these descriptive studies have not elucidated the biochemical events that orchestrate eggshell synthesis, nor the molecular identity of key regulatory proteins required for eggshell melanization. Understanding blood meal digestion and reproductive processes in insect vectors of human disease could lead to the development of selective and safe small molecular inhibitors that may act to reduce the rate of disease transmission. To this end, we have been investigating biochemical processes required for blood meal digestion in the midgut lumen and for complete synthesis of the mosquito eggshell and embryo viability (Isoe et al., 2009; Alabaster et al., 2011; Isoe et al., 2011; Rascon et al., 2011; Isoe et al., 2013; Isoe et al., 2019). We have previously discovered that acetyl CoA carboxylase (ACC) and mosquito lineage-specific eggshell organizing factor 1 (EOF1) are essential for complete eggshell formation and melanization in *A. aegypti* (Alabaster et al., 2011; Isoe et al., 2019). Nearly 100% of eggs oviposited by ACC- and EOF1-deficient females had defective eggshells and non-viable egg phenotypes. However, important blueprints for molecular mechanisms involved in mosquito eggshell formation and melanization are poorly understood and remain to be further explored.

In this study, we identified several additional essential eggshell proteins through RNA interference screening and determined their fecundity, melanization, and viability. We found that Nasrat, Closca, Polehole, and Nudel proteins are predominantly expressed in the ovaries of adult females. Furthermore, these proteins are responsible for maintaining eggshell viability and the melanization process without affecting exochorionic ultrastructures. Our eggshell proteomics identified an additional 168 eggshell structural proteins and enzymes, suggesting mosquito extracellular eggshells are composed of a complex mixture of proteins. These proteins function together to perform the cross-linking, melanization, and sclerotization processes in order to protect the embryo and larva from the environment for long periods of time. These data provide new insights into mosquito ovarian maturation and eggshell synthesis that could lead to key advances in the field of vector control.

## Results

### Systematic RNAi screening of mosquito eggshell proteins

We recently performed RNAi screening for the identification of novel mosquito lineage-specific proteins (Isoe et al., 2019). We discovered that one of the screened genes, EOF1 (eggshell organizing factor 1, AAEL012336), was found to be essential for eggshell formation and melanization in eggs of *A. aegypti* and *A. albopictus*. Since EOF1 was not predicted to contain a secretory signal sequence and was not found from the previously conducted mosquito eggshell proteomic studies (Amenya et al., 2010; Marinotti et al., 2014), we hypothesized that EOF1 is an intracellular protein whose presence is necessary for secretion and activation of unknown eggshell proteins during ovarian follicle maturation. To test this hypothesis, we performed systematic RNAi screening of proteins that have been identified as eggshell proteins from a previous *A. aegypti* proteomic studies (Marinotti et al., 2014) in order to possibly identify potential downstream target eggshell proteins of EOF1. Putative functions and RNAi primers used for each eggshell protein screened are shown in *SI Appendix*, Table S1 and Table S2, respectively. RNAi screening of 34 eggshell proteins showed that six proteins are essential for intact eggshell formation in oviposited eggs (Fig. 1). Among those RNAi positive proteins, Nasrat (AAEL008829) and Closca (AAEL000961) are structural proteins that are known to intimately associate with two other proteins, Polehole and Nudel serine protease, in the perivitelline space (between eggshell and oocyte plasma membrane) of developing ovarian follicles in *D. melanogaster* (Jiménez et al., 2002; Mineo et al., 2017). VectorBase BLAST analysis showed that both Nasrat and Closca shared only approximately 25% of sequence identity between *A. aegypti* and *D. melanogaster*. Three other RNAi positives, DCE2 (AAEL006830), DCE4 (AAEL007096), and DCE5 (AAEL010848), appear to belong to the dopachrome converting enzyme family. Cysteine protease cathepsin L-like (CATL3, AAEL002196) was also found to be essential for proper eggshell formation and melanization in *A. aegypti* mosquitoes (Fig. 1). Because RNAi against other 28 eggshell protein genes showed no abnormal observable egg phenotypes, reduced fecundity, or viability, we focused our effort on these six proteins for further characterization.

**Figure 1.**
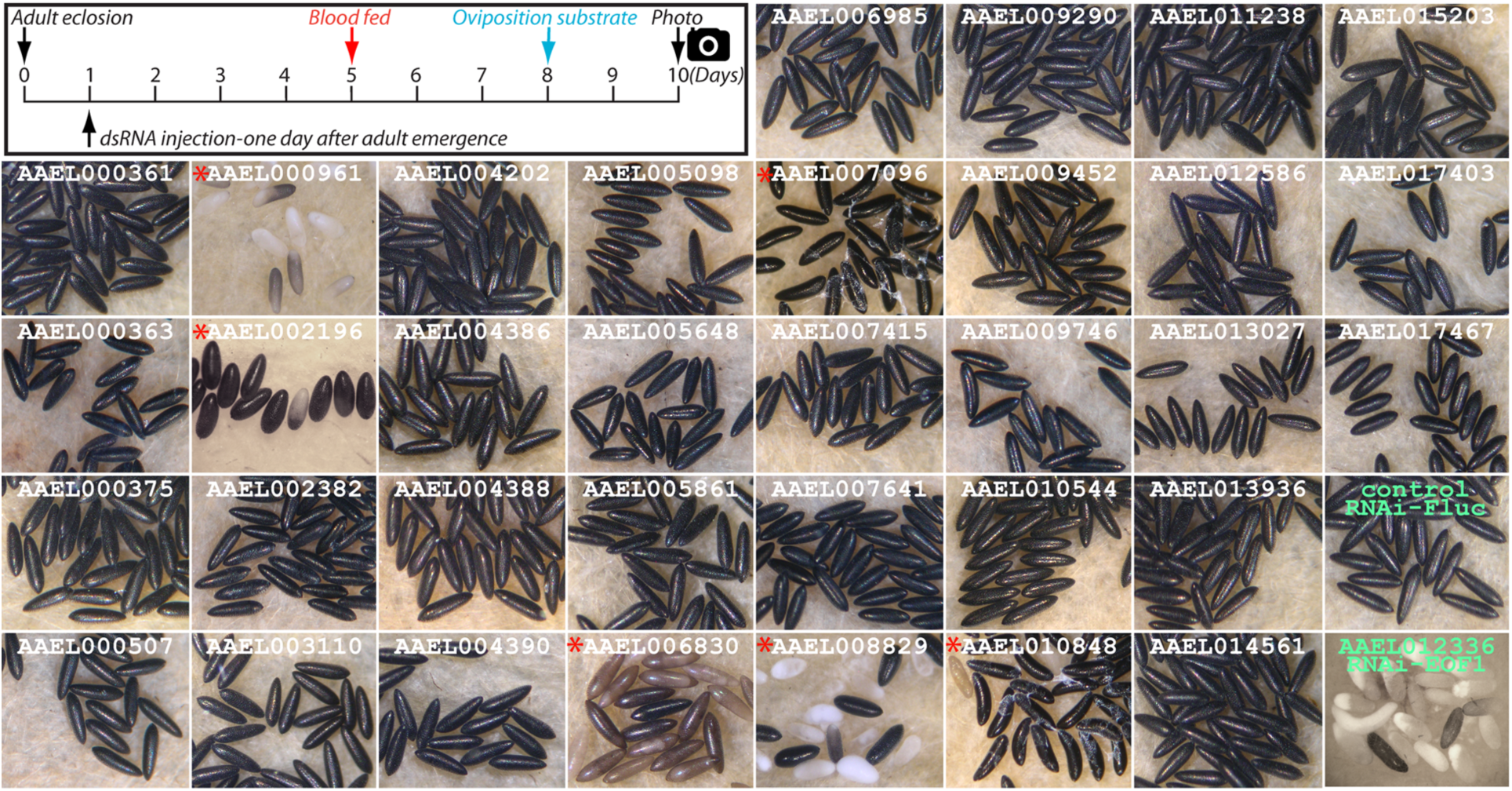
A search for novel essential eggshell proteins in *Aedes aegypti* mosquitoes. Schematic diagram of the experimental time course for dsRNA microinjection, blood feeding, and oviposition in the first gonotrophic cycle shown in the top left corner. Target eggshell proteins were chosen from a proteomic analysis by Marinotti, *et al*., 2014. Representative egg phenotypes associated with RNAi screening of 34 eggshell genes are shown. RNAi against AAEL000961, AAEL002196, AAEL006830, AAEL007096, AAEL008829, and AAEL010848 (as indicated by red asterisks) resulted in defective eggshell phenotypes. Eggs from RNAi-firefly luciferase (Fluc), and RNAi-EOF1 (AAEL012336) are also included. Eggs deposited by ten to twenty RNAi mosquitoes for each respective eggshell protein RNAi knockdown were examined under a light microscope and photographed two days after oviposition. Putative functions and RNAi primers used for each gene are shown in *SI Appendix*, Table S1 and S2, respectively.

### Nasrat, Closca, Polehole, and Nudel are important for eggshell melanization

It has been known that a heterotrimeric protein complex formed by Nasrat, Closca, and Polehole structural eggshell proteins interacts with the Nudel protease during *D. melanogaster* eggshell formation (Jiménez et al., 2002; Mineo et al., 2017). Polehole and Nudel are also highly diverged proteins, sharing 22 and 41% amino acid sequence identity between mosquitoes and fruit flies. Although Polehole and Nudel were not identified as eggshell proteins in the previous mosquito proteomic studies (Amenya et al., 2010; Marinotti et al., 2014), we hypothesized that both proteins may also play important roles in eggshell formation and melanization in *A. aegypti* mosquitoes. We performed RNAi-mediated knockdown studies and observed melanization defective egg phenotypes associated with reduced Polehole and Nudel expression (Fig. 2A). Motivated by the discovery that all four closely associated proteins play essential roles in eggshell melanization, we further characterized reproductive phenotypes, including fecundity, egg melanization, and viability, in response to RNAi-mediated knockdown of these four proteins in the first gonotrophic cycle. First, we examined the effect of RNAi on fecundity by counting eggs oviposited from dsRNA microinjected female mosquitoes. As shown in Fig. 2B, there are no significant differences in the number of eggs oviposited among different RNA deficient mosquitoes compared to the control (*SI Appendix*, Table S3), suggesting that these four proteins play no significant roles in blood meal digestion in midgut lumen, vitellogenin yolk protein synthesis in fat body, and the uptake by primary follicles. Conversely, RNAi silencing of these genes had a significant impact on egg melanization and viability. We observed that nearly 99% of eggs laid by mosquitoes in the control group were melanized. Comparatively, 79 to 84% of eggs deposited by Nasrat, Closca, and Polehole deficient mosquitoes did not undergo complete melanization (Fig. 2C, *SI Appendix*, Table S3). We also observed that females exposed to RNAi-Nudel exhibited the most adverse effect on eggshell melanization. Only 1% of their eggshells completed the melanization process, further indicating that these four proteins may be particularly involved in the formation and melanization of eggshells. Additionally, egg viability was severely impacted by RNAi against these four proteins. Nearly 100% of the eggs deposited by Nudel deficient mosquitoes did not hatch viable larvae and immediately deflated when they were dried. Furthermore, RNAi against Nasrat, Closca, and Polehole resulted in about 10% egg viability (Fig. 2D, *SI Appendix*, Table S3), indicating that complete eggshell melanization may correlate with egg viability, and that these four proteins may likely function together to play an essential role in a molecular and biochemical pathway leading to mosquito eggshell melanization. Quantitative real-time PCR (qPCR) using a single mosquito analysis with gene-specific primers (*SI Appendix*, Table S4) confirmed that the reproductive phenotypes associated with RNAi against Nasrat, Closca, Polehole, and Nudel are likely due to the reduced levels of the corresponding endogenous mRNA in the ovaries (*SI Appendix*, Fig. S1).

**Figure 2.**
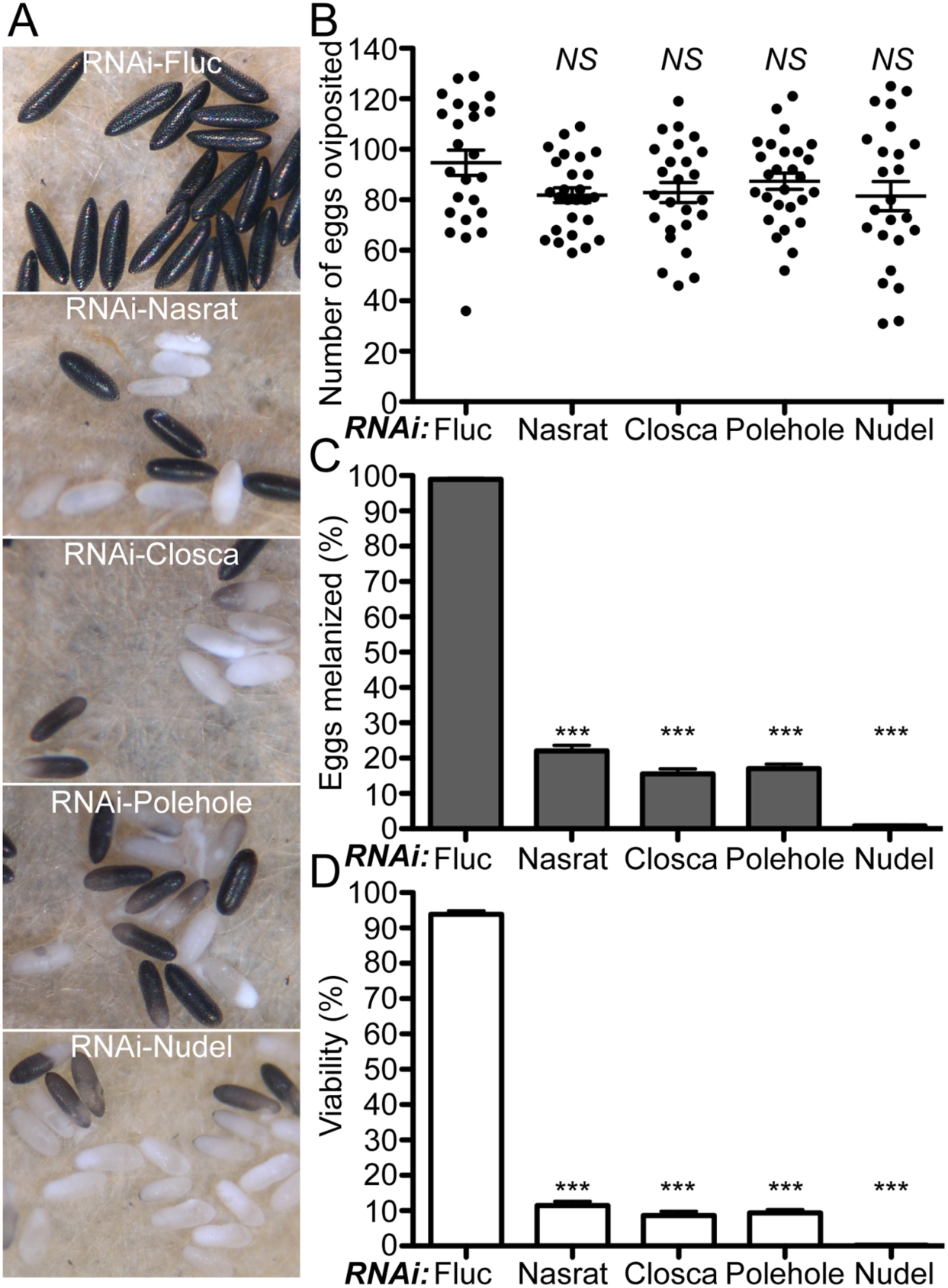
Reproductive phenotypes associated with RNAi against Nasrat, Closca, Polehole, and Nudel in *Aedes aegypti* mosquitoes. (**A**) Representative eggs are shown from mosquitoes microinjected with dsRNA against Fluc control, Nasrat, Closca, Polehole, and Nudel. (**B**) An effect of RNAi on fecundity was studied during the first gonotrophic cycle. Each dot represents the number of eggs oviposited by an individual mosquito. (**C**) Melanization of these eggs was examined under a light microscope and a melanization percentage was determined. An effect of RNAi on viability of eggs was examined by hatching eggs one week after oviposition. Each bar corresponds to egg viability from 15 individual mosquitoes from three independent cohorts. Vectorbase ID: Nasrat (AAEL008829), Closca (AAEL000961), Polehole (AAEL022628), and Nudel (AAEL016971). The mean ± SEM are shown as horizontal lines. Statistical significance is represented by stars above each column (unpaired student’s t test; *** *P* < 0.001, *NS* not significant). A detailed phenotypic analysis is shown in *SI Appendix*, Table S3. Primers used are shown in *SI Appendix*, Table S4.

Next, we determined developmental- and tissue-specific expression patterns of Nasrat, Closca, Polehole, and Nudel during the first gonotrophic cycle in *A. aegypti* by qPCR. Similar adult female- and ovary-predominant expression patterns were observed for Nasrat, Closca, and Polehole. We observed 50 to 200-fold more transcripts in ovaries at 24 h PBM from these three genes compared to the sugar-fed fat body samples. The levels of mRNA encoding for these three eggshell structural proteins in the female reproductive organs increased only slightly in response to blood feeding (Fig. 3A, 3B, and 3C). On the other hand, we found a high level of Nudel serine protease mRNA isolated from the whole body of 3-day old adult male mosquitoes (Fig. 3D), while 200-fold up-regulation of Nudel mRNA expression was detected in ovaries in response to blood feeding. The expression level peaked transiently around 36 hours PBM before dropping precipitously at the end of the primary follicle maturation phase (48 hours PBM, Fig. 3D). Thus, these four proteins may likely play specific roles in the late ovarian follicle development in blood fed *A. aegypti* female mosquitoes, as well as in the processes of eggshell formation and melanization as shown in Fig. 2A. We hypothesized that performing microinjection of dsRNA against these four proteins at specific time points may be important to the knockdown of corresponding mRNAs and may also play a role in inducing detrimental phenotypes of the eggs. Our results show that dsRNA microinjection against Nasrat, Closca, and Polehole immediately after blood feeding did not result in defective egg phenotypes, suggesting that these proteins pre-exist prior to blood feeding, and that they may still be sufficient to function during eggshell formation and melanization (Fig. 4A and 4B and *SI Appendix*, Table S5). In contrast, although fecundity was not affected when dsRNA was microinjected immediately after blood feeding (Fig. 4C), egg melanization and viability were strongly affected in RNAi-Nudel females (Fig. 4D and 4E), indicating that dsRNA microinjection against Nudel at the time of blood feeding may still be capable of silencing its mRNA. This is likely due to the fact that Nudel mRNA transcription is tightly and highly regulated in response to blood feeding, and also because Nudel may function in the late stage of eggshell formation. We recently demonstrated that RNAi-mediated EOF1 depletion resulted in the formation of a defective eggshell in the first three gonotrophic cycles after a single dsRNA injection (Isoe et al., 2019). Thus, we tested whether an effect of RNAi-knockdown of Nudel remains during the development of secondary follicles. Unlike the first gonotrophic cycle, defective phenotypes were no longer observed in eggs deposited by RNAi-Nudel females in the second gonotrophic cycle, and aspects of egg melanization and viability were not significantly affected (Fig. 5 and *SI Appendix*, Table S6). Similarly, abnormal egg phenotypes were not observed in mosquitoes with RNAi against Nasrat, Closca, and Polehole in the second gonotrophic cycle when mosquitoes were microinjected with dsRNA one day after adult eclosion. Because all four proteins may predominantly be expressed and secreted by the follicular epithelial cells, which surround a mature oocyte and undergo a shedding process prior to the departure of the mature primary follicles from an ovariole into an inner oviduct, the secondary follicles and their associated immature state of follicular epithelial cells may not be adversely affected when dsRNA microinjection occurs during the first gonotrophic cycle. Taken together, the Nudel serine protease may interact with Nasrat, Closca, and Polehole structural proteins in the eggshell during the late follicle development stage. The consequence of this interaction may play key roles in the formation and melanization of *A. aegypti* mosquito eggshells.

**Figure 3.**
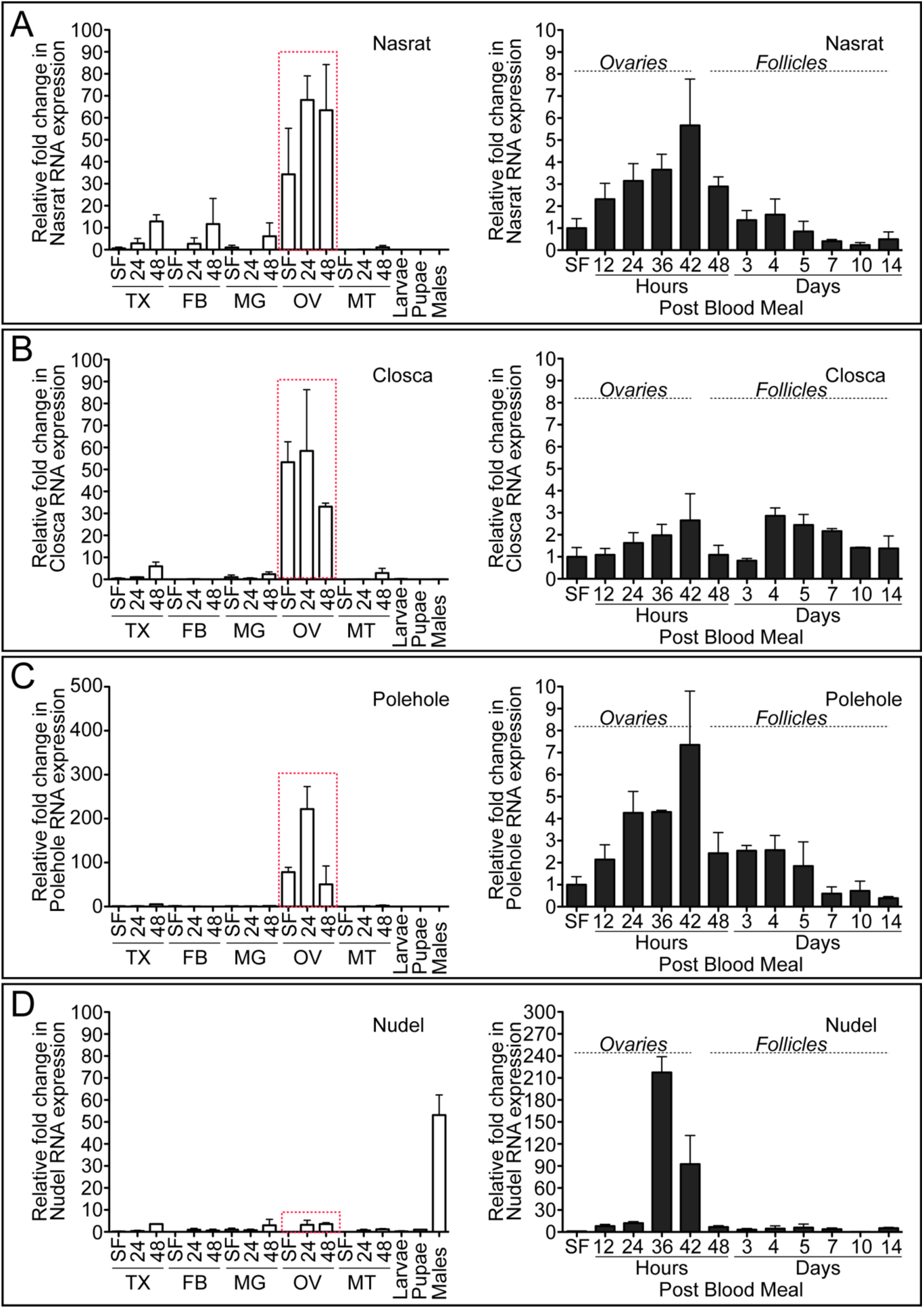
Expression level of Nasrat, Closca, Polehole, and Nudel during the first gonotrophic cycle in *Aedes aegypti* mosquitoes. Tissue-specific and developmental expression pattern of Nasrat (**A**), Closca (**B**), Polehole (**C**), and Nudel (**D**) during the first gonotrophic cycle was determined. The gene expression was analyzed by qPCR using cDNAs prepared from various tissues. Tissues included are thorax (TX), fat body (FB), midgut (MG), ovary (OV), and Malphigian tubules (MT) in sugar-fed only (SF), 24, and 48 hours post blood meal (PBM), as well as larvae (Lv), pupae (Pp), and adult males (M). The expression in fat body at SF was set to 1.0. The tissue expression study shown in the left-hand plots demonstrates that Nasrat, Closca, and Polehole are predominantly expressed in ovaries (highlighted with a red dotted rectangle), while Nudel transcripts are significantly observed in whole body male mosquitoes. A detailed gene expression study in mosquito ovaries or follicles in the first gonotrophic cycle was also analyzed by qPCR. In the right-hand plots, samples from SF to 36 hours PBM include entire ovaries, whereas those from 48 hours to 14 days PBM include only primary follicles isolated from ovaries. Expression in ovaries at SF was set to 1.0. The expression levels for Nasrat, Closca, Polehole, and Nudel were normalized to S7 ribosomal protein transcript levels in the same cDNA samples. Data were collected from three different mosquito cohorts. The mean ± SEM are shown as horizontal lines. Vectorbase ID: Nasrat (AAEL008829), Closca (AAEL000961), Polehole (AAEL022628), and Nudel (AAEL016971). qPCR Primers used are shown in *SI Appendix*, Table S4.

**Figure 4.**
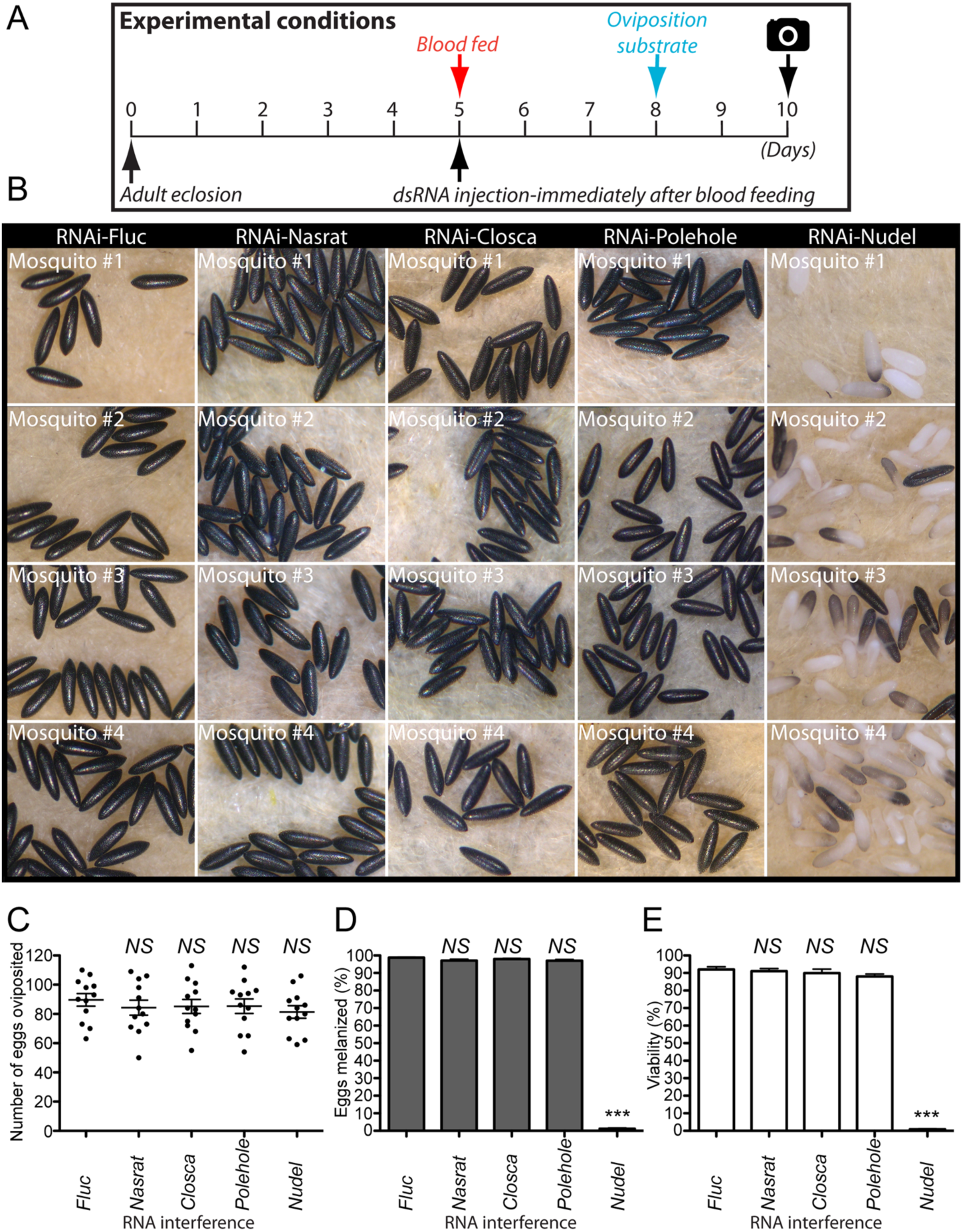
Reproductive phenotypes associated with selected RNAi in response to dsRNA injection immediately after blood feeding. (**A**) Schematic diagram of experimental time course for dsRNA microinjection, blood feeding, and oviposition in the first gonotrophic cycles. Mosquitoes were microinjected with dsRNA immediately after blood feeding. (**B**) Representative eggs shown are from four different mosquitoes either microinjected with dsRNA-Fluc, -Nasrat, -Closca, -Polehole, or -Nudel. Female mosquitoes microinjected with dsRNA-Nasrat, -Closca, or -Polehole laid eggs that showed no difference in fecundity (**C**), melanization (**D**), and viability (**E**) compared to RNAi-Fluc control mosquitoes. Females microinjected with dsRNA-Nudel immediately after blood feeding were adversely affected in terms of eggshell melanization and viability. The effect of selected RNAi on *Ae. aegypti* fecundity was examined by counting the number of eggs laid by each individual female. Each dot represents the number of eggs oviposited by an individual mosquito (**C**). Melanization of these eggs was examined under a light microscope (**D**), and their viability was determined by hatching them one week after oviposition (**E**). Each bar corresponds to egg viability from 15 individual mosquitoes from three groups. The mean ± SEM are shown as horizontal lines. Statistical significance is represented by stars above each column (unpaired student’s t test; *** *P* < 0.001, *NS* not significant). Vectorbase ID: Nasrat (AAEL008829), Closca (AAEL000961), Polehole (AAEL022628), and Nudel (AAEL016971). A detailed phenotypic analysis is shown in *SI Appendix*, Table S5.

**Figure 5.**
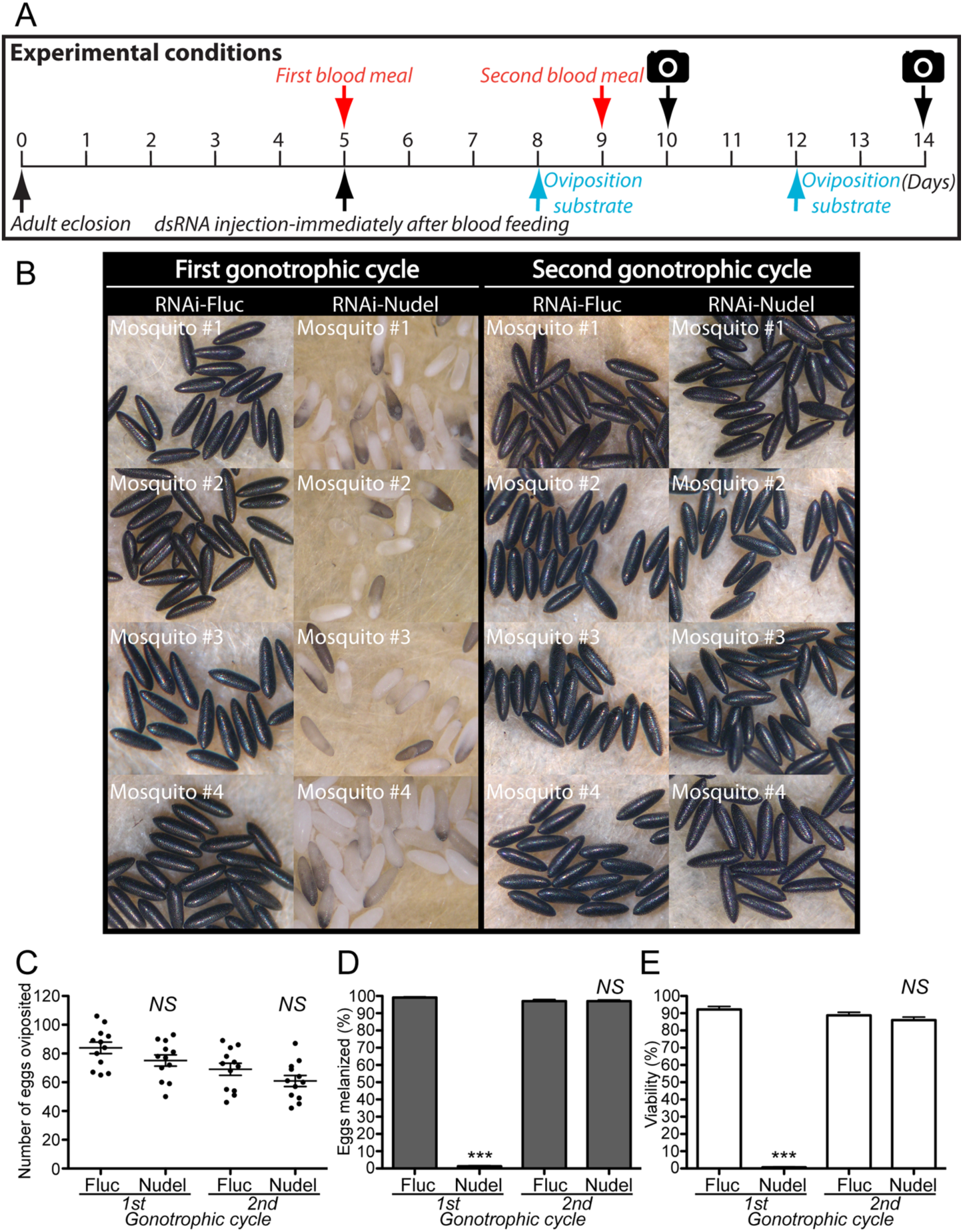
Eggs from the second gonotrophic cycle are not affected by dsRNA-Nudel when microinjected prior or during the blood feeding stage. **A**) Schematic diagram of experimental time course for dsRNA microinjection, blood feeding, and oviposition in the first and second gonotrophic cycles. **B)** Representative eggs are shown from mosquitoes microinjected with dsRNA-Fluc and dsRNA-Nudel. An effect of RNAi-Nudel and RNAi-Fluc control on fecundity (**C**), melanization (**D**), and viability (**E**) was examined during the first and second gonotrophic cycles. Each dot represents the number of eggs oviposited by an individual mosquito (**C**). Melanization of these eggs were examined under a light microscope (**D**), and their viability was determined by hatching them one week after oviposition (**E**). Each bar corresponds to egg viability from 15 individual mosquitoes from three groups. The mean ± SEM are shown as horizontal lines. Statistical significance is represented by stars above each column (unpaired student’s t test; *** *P* < 0.001, *NS* not significant). Vectorbase ID: Nudel (AAEL016971). A detailed phenotypic analysis is shown in *SI Appendix*, Table S6.

### *In vitro* mosquito follicle melanization and permeability assays

Since RNAi-Nudel *A. aegypti* female mosquitoes laid melanization defective eggs, which led to embryonic death (Fig. 2, Video S1 and S2), we hypothesized that activation of the Nudel serine protease in wild type mosquitoes may be tightly regulated temporally and spatially in the perivitelline space. Additionally, we hypothesized that Nudel may likely be inactive prior to oviposition, and that it is immediately activated upon the oviposition event that allows Nudel to proteolytically regulate enzymes involved in melanization, sclerotization, and cross-linking of eggshells. To test our hypothesis, we developed *in vitro* mosquito follicle melanization experiments to determine the time required for melanization of primary follicles isolated from RNAi-mosquitoes. Briefly, ovaries were dissected from wild type or dsRNA injected mosquitoes at 96 h post blood meal. The mature follicles were separated from the ovaries, transferred to oviposition paper wetted with water, and photographed periodically to determine the amount of time required for eggshell melanization. Follicles isolated from RNAi-Fluc controls initiated melanization approximately 90 min after ovary dissection and completed the process approximately three hours after ovary dissection (Fig. 6A). In contrast, the majority of follicles isolated from mosquitoes treated with RNAi against Nasrat, Closca, Polehole, and Nudel did not undergo melanizataion within the three-hour experimental period and never underwent further melanization even 24 hours after the experiments were initiated. These *in vitro* mosquito follicle melanization results suggest that the Nudel serine protease-mediated eggshell melanization may be essential in *A. aegypti* mosquitoes (Fig. 6A and B, and *SI Appendix*, Table S7). Next, to test whether active protease(s) are necessary during eggshell melanization, dissected matured primary follicles were soaked *in vitro* with protease inhibitors (PI, cOmplete™ Mini Protease Inhibitor Cocktail, Roche). Ovaries from female mosquitoes at 96 hours PBM were dissected and placed in water containing PI at different time points after ovary dissection. Individual follicles were then separated, transferred onto oviposition paper, continuously soaked with water containing the PI, and monitored for eggshell melanization. The control follicles were soaked in water without PI. We observed that PI completely inhibited melanization of wild type mature follicles when treated during dissection (Fig. 6C and 6D). On the contrary, the cohort follicles treated with PI at 10 or 20 min after dissection were able to undergo normal eggshell melanization, which was similar to wild type control follicles soaked in water without PI. Thus, these studies with PI suggest that some eggshell proteins may be proteolytically processed immediately after exposure to water in order to activate enzymes involved in eggshell melanization. Next, we treated wild type primary follicles with phenylmethylsulfonyl fluoride (PMSF), which is known to inhibit certain serine proteases. Interestingly, melanization was not blocked when follicles were treated at dissection with PMSF (Fig. 6C). Thus, unknown proteases present in eggshells are essential and activated immediately after oviposition in order to regulate enzymes necessary for eggshell melanization, sclerotization, and cross-linking in *Ae. aegypti*.

**Figure 6.**
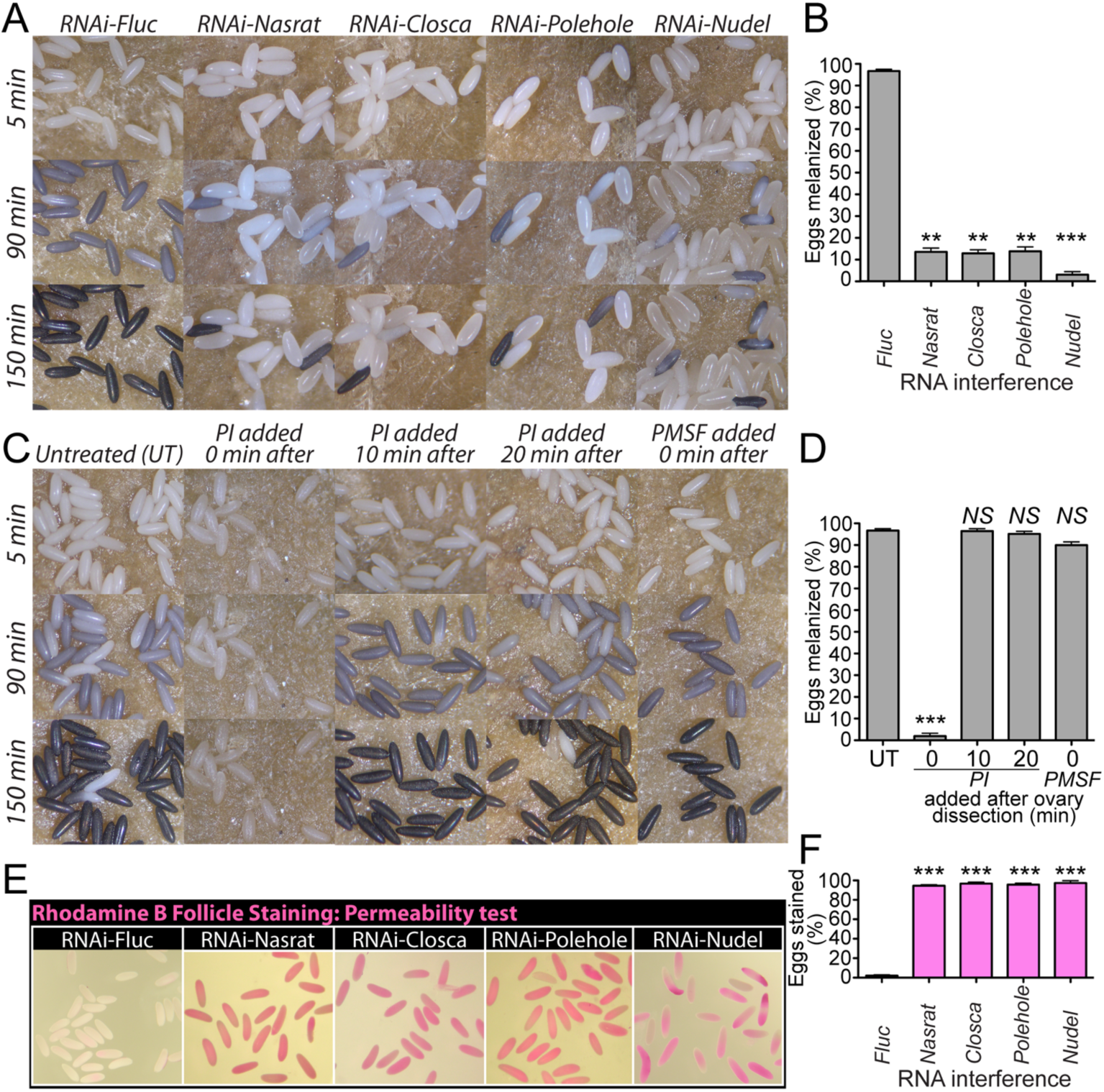
Nasrat, Closca, Polehole, and Nudel are essential for eggshell melanization and oocyte membrane permeability. (**A**) An *in vitro* follicle melanization assay was conducted using follicles isolated from RNAi mosquitoes at 96 hours PBM. The timing of dsRNA microinjection and blood feeding schedule are identical to those shown in Figure 1. The follicles were photographed 5, 90 and 150 min after follicle dissection. (**B**) Each bar corresponds to mean egg melanization (%) from five individual mosquitoes. The mean ± SEM are shown as horizontal lines. Statistical significance is represented by stars above each column (unpaired student’s t test; *** *P* < 0.001, ** *P* < 0.01). (**C**) An *in vitro* follicle melanization assay was performed using a protease inhibitor cocktail (PI) to determine the role of proteases on eggshell melanization. The follicles were photographed 5, 90 and 150 min after follicle dissection. Follicles were incubated with PI (1X) at 0, 10, or 20 minutes after follicle dissection. Follicles were also incubated with PMSF (10 mM) immediately after dissection. (**D**) Each bar corresponds to mean egg melanization (%) from five individual mosquitoes. The mean ± SEM are shown as horizontal lines. Statistical significance is represented by stars above each column (unpaired student’s t test; *** *P* < 0.001, *NS* not significant). (**E**) The follicle permeability assays were conducted using a Rhodamine B marker. A Rhodamine B cellular uptake was observed in primary follicles possibly due to a defective eggshell and oocyte plasma membrane. Vectorbase ID: Nasrat (AAEL008829), Closca (AAEL000961), Polehole (AAEL022628), and Nudel (AAEL016971). A detailed phenotypic analysis is shown in *SI Appendix*, Table S7 and Table S8.

Next, we determined whether water permeability of the ovarian primary follicles is affected due to defective eggshells by staining the dissected follicles from RNAi mosquitoes at 96 hours PBM with a rhodamine B marker. Rhodamine B permeability assays have an advantage that they can quickly assess whether follicles within the ovaries contain defective eggshells prior to oviposition. The representative images of follicles from RNAi are shown in Fig. 6E and 6F. Follicles from RNAi-Fluc mosquitoes were only slightly stained by rhodamine B, while increased permeability and uptake of rhodamine B through an oocyte plasma membrane in mosquitoes with RNAi against Nasrat, Closca, Polehole, and Nudel was observed (Fig. 6E, *SI Appendix*, Table S8), indicating that defective eggshells caused by a loss of these proteins may not properly regulate water permeability into oocytes when eggs are oviposited. Taken together, we speculate that these four eggshell proteins play a significant role in important eggshell formation processes and thus egg viability.

### Phenotypic characterization of eggs deposited by RNAi-DCEs and -CATL3 mosquitoes

In addition to four proteins studied above, RNAi screening identified four mosquito eggshell proteins, DCE2, DCE4, DCE5, and CATL3, that are necessary to form intact eggshells (Fig. 1). We performed RNAi-mediated knockdown studies to further characterize reproductive phenotypes, including fecundity, egg melanization, and viability in the first gonotrophic cycle. qPCR using gene-specific primers (*SI Appendix*, Table S4) validated that RNAi against DCE2, DCE4, DCE5, and CATL3 had significant effects on the reduced level of the corresponding endogenous mRNA in the ovaries (*SI Appendix*, Fig. S2). Representative egg phenotypes from RNAi treated mosquitoes are shown in Fig. 7A. RNAi-mediated loss of either DCE2, DCE4, DCE5, or CATL3 did not significantly affect fecundity, compared to the RNAi-Fluc control (Fig. 7B, *SI Appendix*, Table S9). RNAi-silencing of DCE2 and CATL3 had differential effects on eggshell melanization. Approximately 96% of total eggs from RNAi-DCE2 females were only partially melanized, while 43% of eggs from RNAi-CATL3 females were totally non-melanizaed (Fig. 7C, *SI Appendix*, Table S9). On the other hand, nearly all eggs deposited by RNAi-DCE4 and -DCE5 mosquitoes were melanized normally with readily observable defective eggshell surface phenotypes, suggesting that DCE4 and DCE5 eggshell enzymes may not be involved in endochorion melanization and play important roles in exochorion formation. Approximately 95% of eggs from RNAi-DCE2 and – CATL3 females did not contain viable offspring, while nearly 90% of laid by RNAi-DCE4 and -DCE5 mosquitoes had viable first instar larvae.

**Figure 7.**
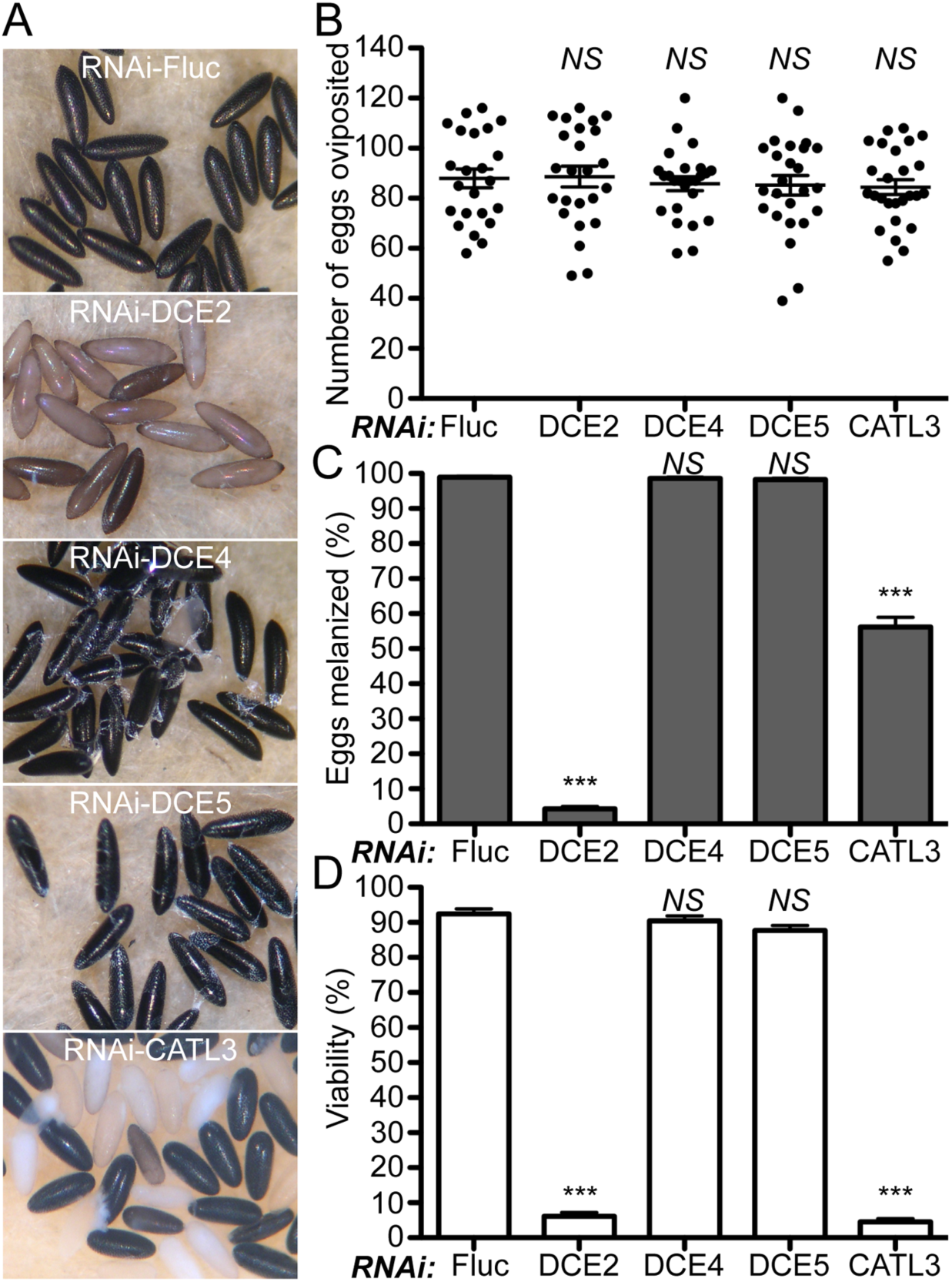
Reproductive phenotypes associated with RNAi against DCE2, DCE4, DCE5, and CATL3 in *Aedes aegypti* mosquitoes. (**A**) Representative eggs are shown from mosquitoes microinjected with dsRNA against Fluc control, DCE2, DCE4, DCE5, and CATL3. (**B**) An effect of RNAi on fecundity was studied during the first gonotrophic cycle. Each dot represents the number of eggs oviposited by an individual mosquito. (**C**) Melanization of these eggs was examined under a light microscope and a melanization percentage was determined. (**D**) An effect of RNAi on viability of eggs was examined by hatching eggs one week after oviposition. Each bar corresponds to egg viability from 15 individual mosquitoes from three independent cohorts. Vectorbase ID: DCE2 (AAEL006830), DCE4 (AAEL007096), DCE5 (AAEL010848), and CATL3 (AAEL002196). The mean ± SEM are shown as horizontal lines. Statistical significance is represented by stars above each column (unpaired student’s t test; *** *P* < 0.001, *NS* not significant). A detailed phenotypic analysis is shown in *SI Appendix*, Table S9. Primers used are shown in *SI Appendix*, Table S4.

Next, we examined developmental- and tissue-specific expression patterns of DCE2, DCE4, DCE5, or CATL3 during the first gonotrophic cycle in *A. aegypti* by qPCR. We observed that all four genes exhibited clear adult female- and ovary-specific expression patterns (Fig. 8). Only a 5-fold increase in DCE2 ovarian mRNA was observed in response to blood feeding (Fig. 8A). DCE4 and DCE5 mRNA expression patterns were similar to each other and highly induced in ovaries after blood feeding, peaking at around 42 hours PBM (Fig. 8B and 8C). Highly up-regulated CATL3 mRNA transcripts were observed in ovaries between 36 and 42 hours PBM (Fig. 8D). The mRNA transcripts encoding DCE4, DCE5, and CATL3 were no longer observed by 72 hours PBM, suggesting that their expressions may be exclusively present in the follicular epithelial cells that undergo shedding and apoptosis when follicles are ready for oviposition.

**Figure 8.**
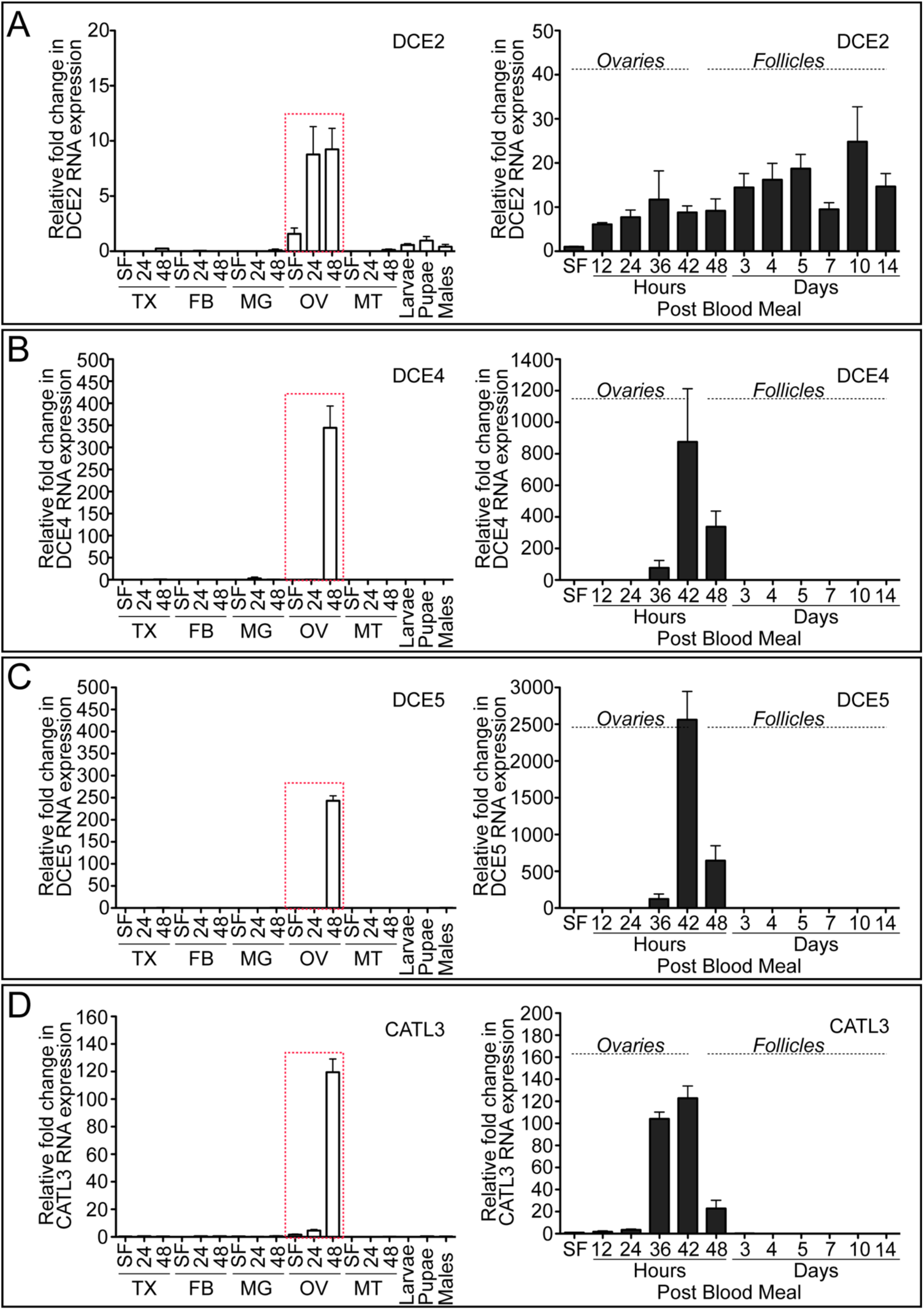
Expression level of DCE2, DCE4, DCE5, and CATL3 during the first gonotrophic cycle in *Aedes aegypti* mosquitoes. Tissue-specific and developmental expression pattern of DCE2 (**A**), DCE4 (**B**), DCE5 (**C**), and CATL3 (**D**) during the first gonotrophic cycle was determined. The gene expression was analyzed by qPCR using cDNAs prepared from various tissues. Tissues included are thorax (TX), fat body (FB), midgut (MG), ovary (OV), and Malphigian tubules (MT) in sugar-fed only (SF), 24, and 48 hours post blood meal (PBM), as well as larvae (Lv), pupae (Pp), and adult males (M). The expression in fat body at SF was set to 1.0. The tissue expression study shown in the left-hand plots demonstrates that DCE2, DCE4, DCE5 and CATL3 are predominantly expressed in ovaries (highlighted with a red dotted rectangle). A detailed gene expression study in mosquito ovaries or follicles in the first gonotrophic cycle was also analyzed by qPCR. In the right-hand plots, samples from SF to 36 hours PBM include entire ovaries, whereas those from 48 hours to 14 days PBM include only primary follicles isolated from ovaries. Expression in ovaries at SF was set to 1.0. The expression levels for DCE2, DCE4, DCE5, and CATL3 were normalized to S7 ribosomal protein transcript levels in the same cDNA samples. DCE4, DCE5 and CATL3 transcripts are significantly up-regulated during late phase ovarian follicle development. Data were collected from three different mosquito cohorts. The mean ± SEM are shown as horizontal lines. Vectorbase ID: DCE2 (AAEL006830), DCE4 (AAEL007096), DCE5 (AAEL010848), and CATL3 (AAEL002196). qPCR Primers used are shown in *SI Appendix*, Table S4.

### Ultrastructural analysis of defective eggshells under a scanning electron microscope

We further characterized the effect of RNAi on topological surface features of the mosquito eggshell in detail using a scanning electron microscope (SEM). SEM images of an entire primary follicle and its detailed surface features from RNAi-Fluc control samples are shown, respectively, for comparison (Fig. 9A and 9B). Ultrastructural analysis of the exochorion outermost layer of the eggshell from *A. aegypti* has been previously carried out under SEM (Pereira et al., 2006; Suman et al., 2011; Bova et al., 2016; Faull and Williams, 2016). As shown in Fig. 9B, the outer chorionic area (OCA), surrounded by the porous exochorionic network (EN), contains a single protruding central tubercle (CT) as well as several minute peripheral tubercles (PTs). We examined eggs that exhibited defective phenotypes during RNAi screening of eggshell proteins (Fig. 1), as well as Nudel, Polehole, and EOF1. Nudel serine protease and Nasrat, Closca, and Polehole structural proteins are eggshell proteins that intimately interact with one another. Among these four proteins, we found that RNAi-Nudel showed the strongest melanization effect on eggs (Fig. 2). However, SEM analysis showed that eggs from RNAi-Nudel exhibited eggshell surface features that were similar to the control (Fig. 9C). Likewise, exochorionic surface topology of eggs from RNAi-Nasrat, -Closca, and -Polehole females did not contain any dramatically altered phenotypes (Fig. 9D-9F), while eggs from these cohorts had defective eggshell melanization phenotypes similar to those observed in Fig. 2. Although RNAi-DCE2 eggs were incompletely melanized (Fig. 1), the SEM analysis found no detectable abnormal surface phenotypes (Fig. 9G). Thus, SEM studies on these five eggshell proteins suggest that they may likely play key roles in melanization of an endochorion layer beneath the exochorion. On the other hand, RNAi against DCE4 and DCE5 resulted in phenotypes in which the outermost exochorion appears to be partially peeled or entirely absent (Fig. 1). The SEM images confirmed that a portion of the EN shrank and peeled away, while CT and PTs were less affected (Fig. 9H and 9I), suggesting that DCE4 and DCE5 enzymes may possibly be responsible for cross-linking EN to the exochorion surface. Eggs deposited by CATL3-deficient mosquitoes are generally more oval-shaped than those from other RNAi mosquitoes (Fig. 1 and Fig. 7), and we found that RNAi-CATL3 also led to deformed EN structures under SEM (Fig. 9J). However, these ultrastructural features observed (Fig. 9B-9J) are dissimilar to eggshell surface characters resulting from RNAi-EOF1 (Fig. 9K, Isoe et al., 2019), where an enlarged OCA contains multiple CTs. Since a loss of EOF1 function by RNAi led to altered exochorion ultrastructural features and defects in endochorion melanization, the mosquito-specific intracellular EOF1 protein may regulate multiple target proteins that are independently responsible for eggshell surface patterning of exochorion layer and melanization of endochorion layer in *A. aegypti* mosquitoes.

**Figure 9.**
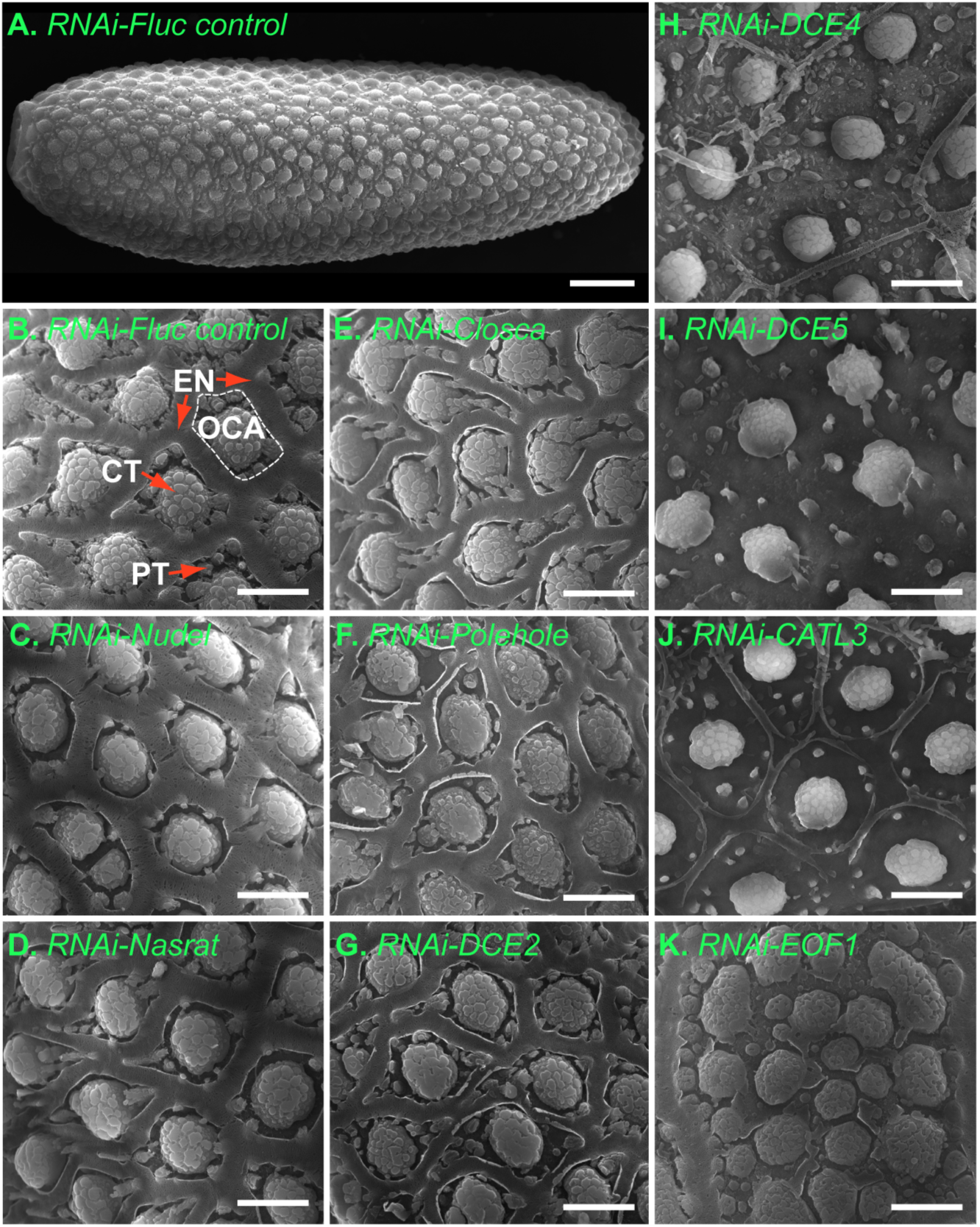
Representative scanning electron micrograph images of eggshell ultrastructural features from RNAi *Aedes aegypti* mosquitoes. **A**) Dorsal view of an entire primary follicle from an RNAi-Fluc mosquito. The image was taken under 600X magnification (scale bars = 50 µm). **B**) Representative dorsal images are shown from RNAi-Fluc control (B), -Nudel (C), -Nasrat (D), – Closca (E), -Polehole (F),DCE2 (G), -DEC4 (H), -DCE5 (I), -CATL3 (J), and -EOF1 (K) mosquitoes. The photos were taken under 6,000X magnification (scale bars = 10 µm). Mosquitoes were injected with dsRNA, as shown in Figure 1. The mature primary follicles were dissected in 1X PBS at 96 hours PBM, immediately fixed with 2.5% EM grade glutaraldehyde at 4°C, post-fixed with 1% osmium tetroxide at room temperature, dehydrated with 100% ethanol, dried with hexamethyldisilazane, and metallized with gold. All dorsal images of eggs were taken using a FEI Inspect-S electron scanning microscope at the W.M. Keck Center for Nano-Scale Imaging, University of Arizona. Vectorbase ID: Nudel (AAEL016971), DCE2 (AAEL006830), DCE4 (AAEL007096), CATL3 (AAEL002196), and EOF1 (AAEL012336). We allowed the cohort of mosquitoes microinjected with these dsRNAs to lay eggs and confirmed similar defective eggshell phenotypes, as shown in Fig.1, Fig 2, and Fig. 7.

### Proteomic analysis on mosquito eggshells

Extracellular eggshell structural proteins and enzymes have been identified in previous mosquito eggshell proteomic analyses (Amenya et al., 2010; Marinotti et al., 2014). Specifically, enzymes including chitinases, chorion peroxidases, prophenoloxidases, dopachrome converting enzymes, laccase-like multicopper oxidase, proteases, and others have been found and recognized for their importance in the eggshell formation. We recently identified a novel mosquito lineage-specific EOF1 protein through RNAi screening. EOF1 plays an upstream regulatory role in the process of eggshell melanization in *A. aegypti* mosquitoes (Isoe et al., 2019). Our SEM analysis of mosquito eggs shows that RNAi against eight proteins and enzymes did not phenocopy the severely altered defective eggshell surface topology observed in RNAi-EOF1 (Fig. 9). Thus, the target eggshell proteins of EOF1 remain to be unknown. To determine whether EOF1 deficiency in mosquitoes leads to modulated levels of the downstream target proteins in eggshells, we performed proteomic analysis of eggshells derived from the primary follicles of RNAi-Fluc control and RNAi-EOF1 mosquitoes. Our current study employed a different approach that used Dounce homogenizers to enrich extracellular eggshells from ovarian follicle cells. Eggshell proteins were subsequently extracted with guanidine hydrochloride. The major steps involved are shown in Fig. 10A-10F, and our current mass spectrometry analysis detected a total of 220 eggshell proteins (Fig. 10G, *SI Appendix*, Table S10).

**Figure 10.**
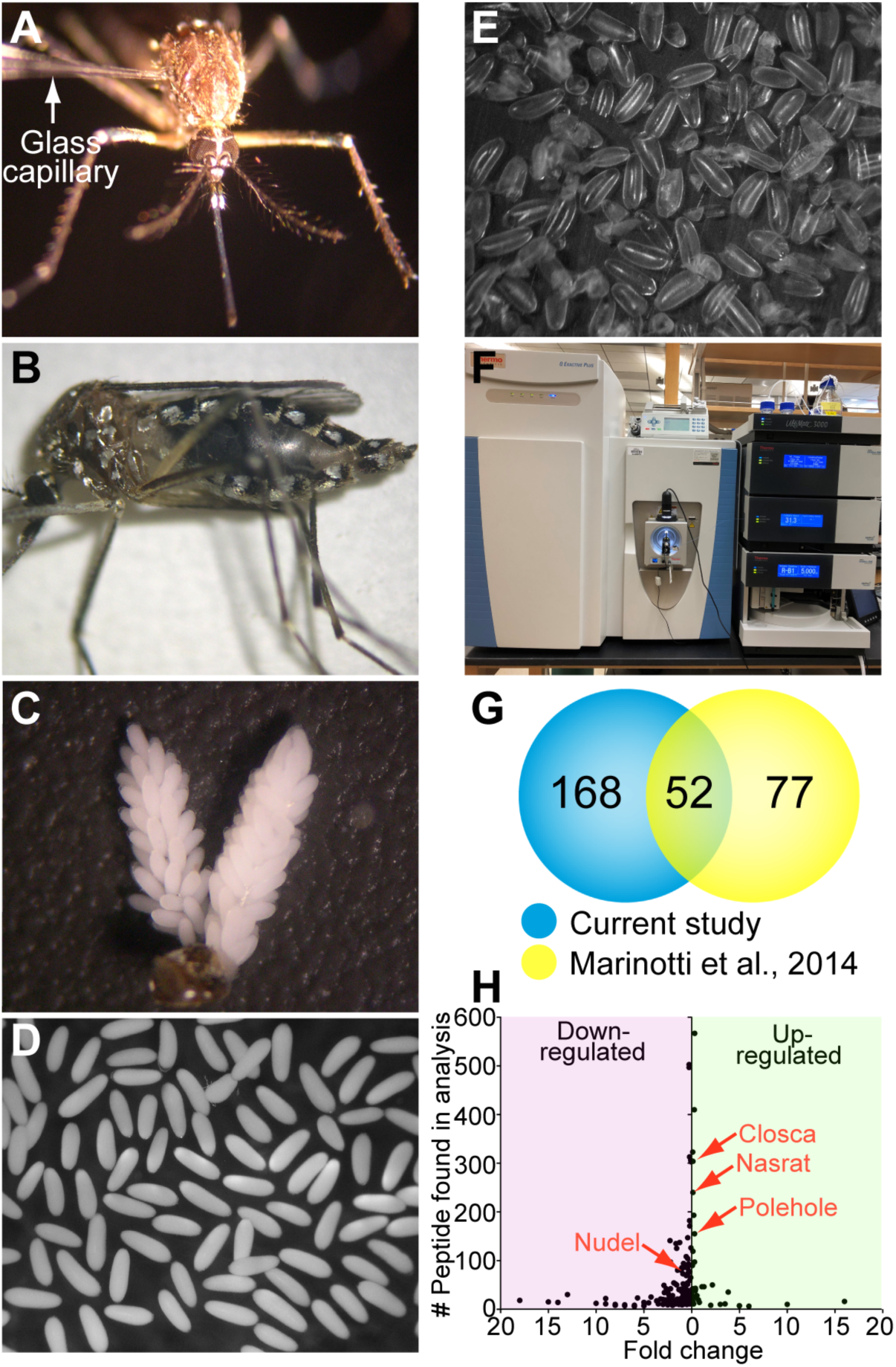
Approaches used for eggshell isolation and eggshell protein identification in *Aedes aegypti*. (**A**) dsRNA samples were microinjected into the mosquito thorax three days prior to blood feeding, as shown in Figure 1. (**B**) dsRNA injected mosquitoes were allowed to feed on blood until fully engorged. (**C**) Ovaries were dissected from dsRNA injected mosquitoes four days post blood meal. (**D**) Primary follicles were purified from dissected ovaries. (**E**) The primary follicles were homogenized using a dounce homogenizer to remove oocyte cytosolic and membrane contents. Eggshells were filtered through a mesh strainer (40 μm) to obtain enriched eggshell. (**F**) Trypsin-digested eggshell proteins were subjected to LC-MS/MS using the Q Exactive™ hybrid quadrupole-Orbitrap mass spectrometer at the University of Arizona Analytical and Biological Mass Spectrometry Facility. (**G**) Venn diagram comparing the number of identified eggshell proteins between the current study and previously published data by Marinotti *et al*., 2014. Only proteins with more than or equal to six peptide hits from four independent proteomic data were included in this study. About 40% of the eggshell proteins from the previous study were found in the current study, while 162 additional proteins were detected in the eggshell. (**H**) Eggshell peptide abundance fold changes in response to RNAi-EOF1 (AAEL012336) are shown in comparison with RNAi-Fluc control. Two independent biological replicates from both RNAi-Fluc and RNAi-EOF1 were used in the proteomic analysis.

Eight putative enzyme and protein functions were further evaluated by comparing from previously published proteomic data (Marinotti et al., 2014) with identified eggshell proteins in this study (Fig. 11). Specifically, extracellular chorion peroxidases are involved in the formation of rigid, insoluble eggs by catalyzing an eggshell protein cross-linking reaction (Han et al., 2000; Li et al., 1996; Li et al., 2004). In our analysis, we found four additional proteins with a chitinase domain, and also discovered two extra chorion peroxidases besides the five that were detected previously (Marinotti et al., 2014). Taken together, the data suggests that the existence of these enzymes may be advantageous to quickly harden eggshells in order to protect fertilized oocytes undergoing embryogenesis. Furthermore, each individual enzyme may have preferred protein substrates and/or localize differently within the eggshell. Similarly, we identified five new putative prophenoloxidases, which may play key roles in the process of eggshell melanization in mosquitoes (Li et al., 1996). Insect dopachrome conversion enzymes (DCE) have been studied in *A. aegypti* (Johnson et al., 2013), *D. melanogaster* (Han et al., 2002), *Tribolium castaneum* (Arakane et al., 2010), and *Musca domestica* (Heinze et al., 2017), since they are responsible for catalyzing melanization and sclerotization reactions. We now know that at least five extracellular DCE are present to form intact eggshells in *A. aegypti* and that they may exhibit specific functions (Fig. 11). We demonstrated this specificity by using RNAi bioassays, which showed that DCE2 is required for melanization (Fig 7A), while DCE4 and DCE5 play a role in the formation of the fine EN structures that define OCA at the eggshell exochorion (Fig. 9H and 9I). Surprisingly, we found 11 new eggshell proteases including Nudel, but we did not detect CATL3, which was found in a previous eggshell proteomic study (Marinotti et al., 2014), and is an essential cysteine protease for egg viability demonstrated in this study (Fig. 7). Additional RNAi knockdown studies against newly identified proteases may reveal their essential functions in mosquito eggshells. The mass spectrometry data from previous studies and our current study have shown that there are a combined 48 different putative odorant-binding proteins with unknown functions in the *A. aegypti* eggshell (Fig. 11, *SI Appendix*, Table S10). The previous study identified seven vitelline membrane proteins, while in the current proteomic study we detected two of those proteins along with a novel protein. Our method for mosquito eggshell isolation failed to detect any putative trypsin inhibitor-like proteins or serpins. Taken together, we discovered 168 new proteins, confirmed the presence of 52 proteins and failed to identify 77 proteins previously identified from the *A. aegypti* eggshell, suggesting that different eggshell isolation and protein extraction methods may be ideally suited for enriching different eggshell proteins. We did not observe a dramatic effect of RNAi-EOF1 on peptide abundance for Nasrat, Closca, Polehole, Nudel, DCE2, DCE4, and DCE5, compared to the RNAi-Fluc control (Fig. 10H, *SI Appendix*, Table S10). It is possible that EOF1 deficiency could result in the mislocalization of these proteins without affecting expression levels, while still leading to the defective phenotypes. Alternatively, other newly identified uncharacterized eggshell proteins (*SI Appendix*, Table S10) may be essential for melanization and cross-linking processes and downstream targets of EOF1.

**Figure 11.**
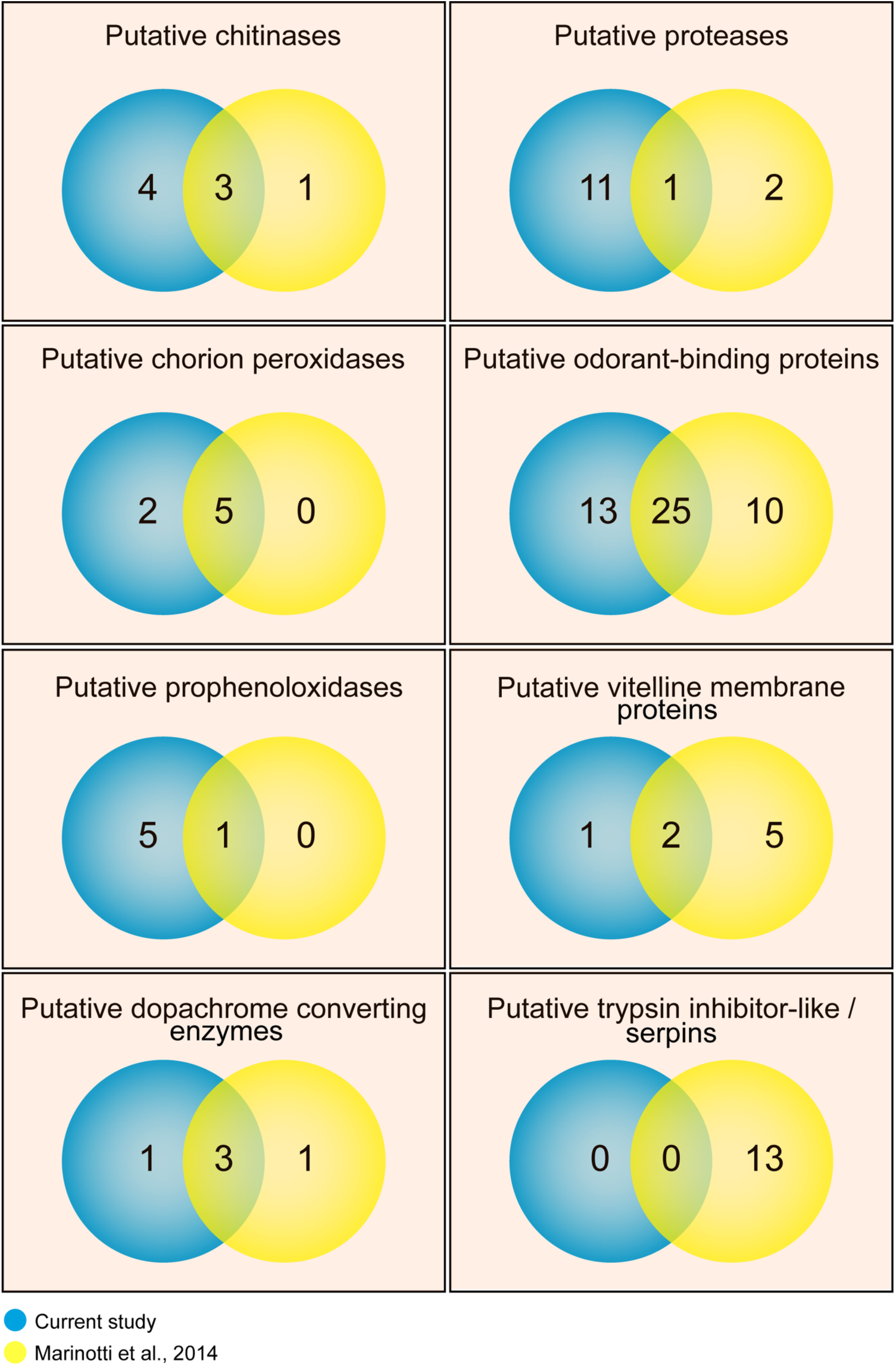
Venn diagram illustrating the identification of proteins in different eggshell proteomic data sets. Eight putative protein functions were evaluated by comparing identified eggshell proteins from this study with previously published proteomic data by Marinotti *et al*., 2014. Proteins with more than or equal to six peptide hits from four independent proteomic data sets were included. Two independent biological replicates from both RNAi-Fluc and RNAi-EOF1 were used in the proteomic analysis.

## Discussion

Zoonotic disease transmission by mosquito vectors is a significant problem throughout the world. One approach to controlling mosquito populations is to disrupt specific molecular processes or antagonize novel metabolic targets required for the production of viable eggs. The eggshell extracellular matrix between follicular epithelial cells and an oocyte is formed in the ovaries, and the melanization and cross-linking occur after oviposition to protect the embryo from the environment. A fully matured *Aedes aegypti* mosquito eggshell can protect the first instar larva for over a year prior to hatching. Here, we identified several essential eggshell proteins and enzymes through RNA interference screening and characterized their associated egg phenotypes in *A. aegypti*.

A proposed model of eggshell formation and melanization in *A. aegypti* mosquitoes is shown in Fig. 12. A primary follicle reaches maturity by up-taking vitellogenin yolk proteins around 48 hours post blood meal (PBM), while a secondary follicle attached to a germarium remains dormant in the resting stage. The extracellular eggshell matrix is sequentially built between the oocyte and follicular epithelial cells and completes its formation by 48 h PBM. SEM images reveal a very complex exochorion (Fig. 9) and endochorion (Fig. 13A and 13B) structure. Mosquito Nasrat, Closca, and Polehole may form a heterotrimeric complex and localize at the perivitelline fluid side of the endochorion, as shown in *D. melanogaster* (Jiménez et al., 2002; Mineo et al., 2017). The Nudel serine protease may be highly accumulated and tethered by the complex in late follicle development. DCE2 may be localized at the endochorion, while DCE4 and DCE5 may be specifically found in the exochorion. Other enzymes responsible for melanization, sclerotization, and cross-linking may likely be present as inactive forms in appropriate locations within the eggshells. Once the primary follicles are ready for ovulation and ovipositon, follicular epithelial cells surrounding the eggshell along with the secondary follicle and germarium undergo shedding processes. These occur in an ovariole, and progress from the posterior to anterior direction (Fig. 13C). The follicles move through inner, lateral, and common oviducts before the fertilized eggs are eventually deposited onto a wet oviposition substrate. Surrounding water is instantaneously and briefly uptaken by the eggs. Nudel and/or CATL3 in eggshells are activated through unknown mechanisms, and they may in turn proteolytically activate enzymes involved in eggshell melanization and cross-linking processes.

**Figure 12.**
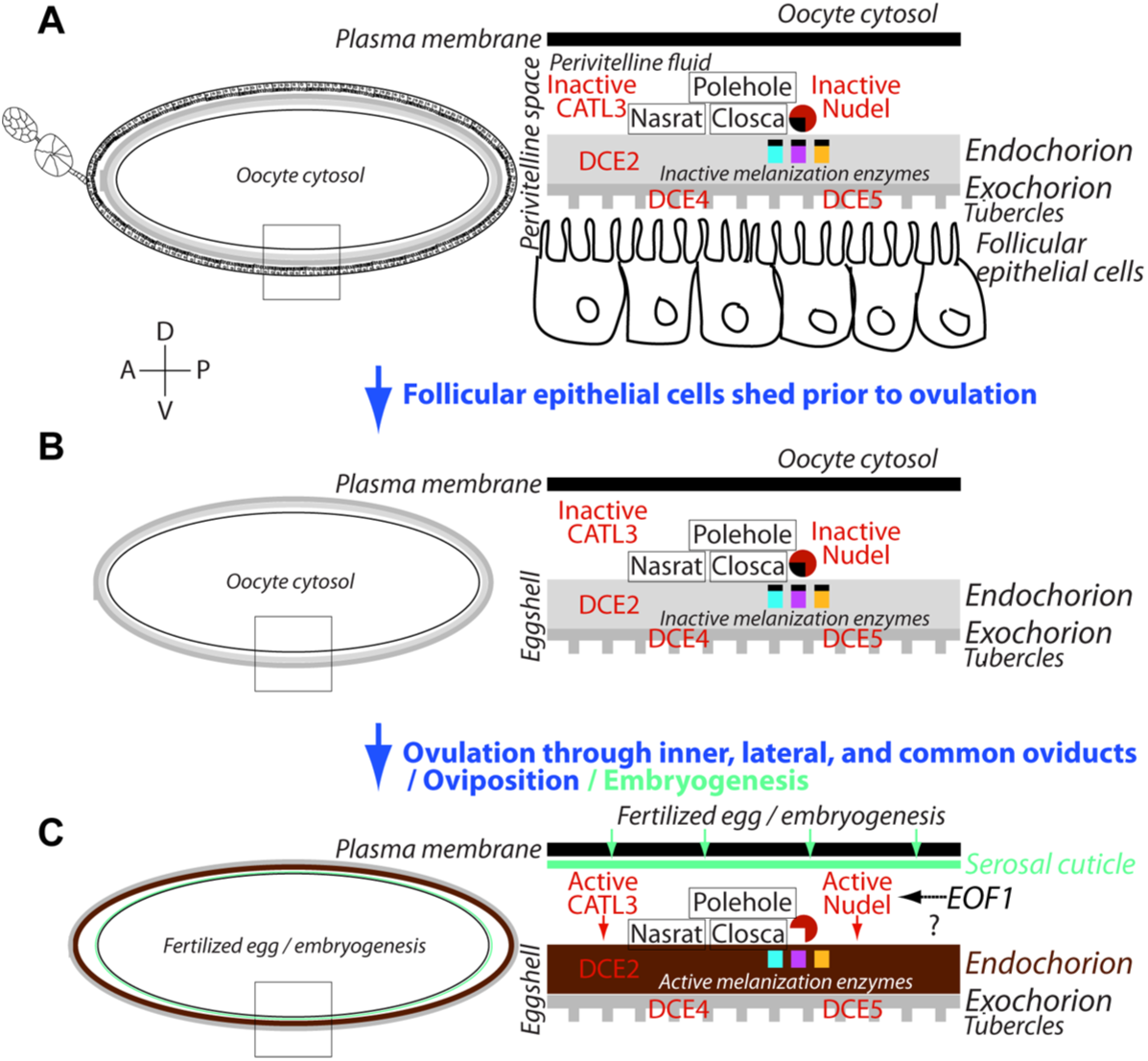
A proposed schematic model of involvement of specific proteins during eggshell formation and melanization in *Aedes aegypti* mosquitoes. (**A**) A primary follicle in the ovariole reaches maturity, and an extracellular eggshell matrix completely forms between oocyte and follicular epithelial cells around 48 hours post blood meal (PBM). Enzymes involved in eggshell melanization and cross-linking may likely be inactive during this stage. (**B**) As the follicles initiate migration into an inner oviduct, the surrounding follicular epithelium sheds with the secondary follicle and germarium in the ovariole. The enzymes involved in eggshell melanization and cross-linking may still be their inactive forms within the inner, lateral, and common oviducts. (**C**) Once the eggs are deposited onto a dump substrate, eggs briefly uptake surrounding water and activate the melanization and cross-linking enzymes through the activity of Nudel and/or CATL3. Vectorbase ID: Nasrat (AAEL008829), Closca (AAEL000961), Polehole (AAEL022628), and Nudel (AAEL016971), DCE2 (AAEL006830), DCE4 (AAEL007096), DCE5 (AAEL010848), and CATL3 (AAEL002196).

**Figure 13.**
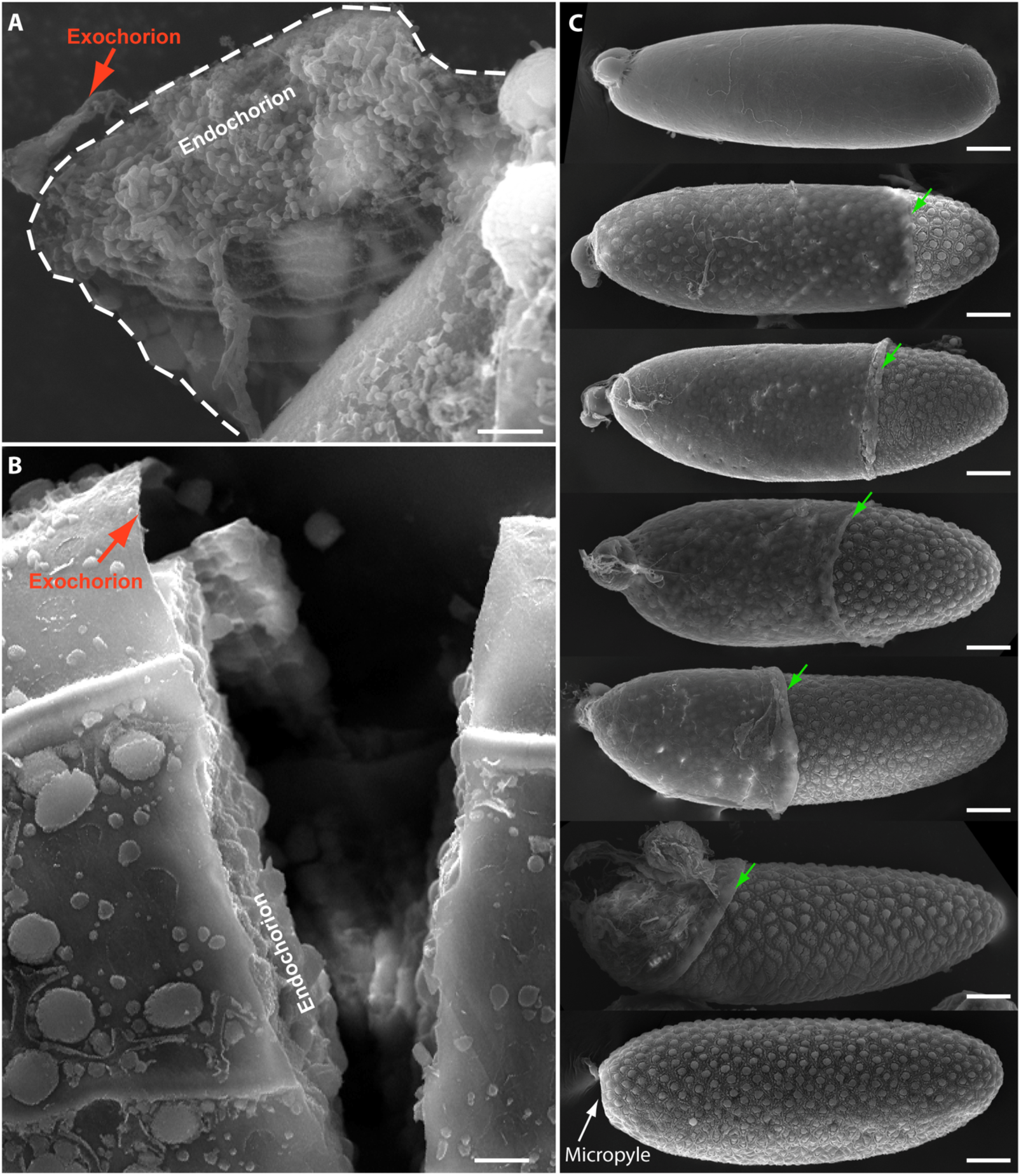
Scanning electron micrograph images of late follicle development in *Aedes aegypti*. (**A**) SEM image shows a peeled area of exochorion and endochorion, exposing the complex internal eggshell structures. (**B**) SEM image shows a vertical breakage of mosquito egg, exposing a very thin exochorion and the complex endochorion eggshell structures. (**C**) Scanning electron micrograph images of individual follicles during late-oogenesis, showing gradual shedding of the follicular epithelial cell layer from the posterior to anterior direction. Programmed shedding of follicular epithelial cells, secondary follicles, and germarium from the primary follicle in the ovariole occurs during late oocyte development prior to ovipositon. Green arrows indicate a position of a leading edge of the follicular epithelial cell layer. Follicles were derived from wild type mosquitoes. Follicle samples were prepared as described in Fig. 9 legend and in the materials and methods. Images were obtained using an FEI Inspect-S electron scanning microscope at the W.M. Keck Center for Nano-Scale Imaging, University of Arizona. The photos were taken under 6,000X magnification (**A**; scale bar 5 µm), under 1,200X magnification (**B**; scale bar 10 µm,), and under 500X magnification (**C**; scale bars 50 µm).

In this study, we have identified several additional essential eggshell proteins through RNA interference screening and determined their egg phenotypes including fecundity, melanization, and viability. All eight characterized eggshell proteins were required for intact eggshell formation. We found that Nasrat, Closca, Polehole, Nudel, DEC4, DCE5, and CATL3 proteins are predominantly expressed in the ovaries of adult females. Furthermore, Nasrat, Closca, Polehole, Nudel, and DCE2, are responsible for maintaining the integrity of eggshell viability and the melanization process and do not affect exochorionic ultrastructures. On the other hand, the functions of DEC4, DCE5, and CATL3 are coupled to the dynamic exochorionic surface topology of mosquito eggs. However, their surface topologies in response to RNAi did not phenocopy the effect of RNAi-EOF1. Thus, it remains unclear how EOF1 influences eggshell formation and melanization. The use of proteomic analysis of eggshell proteins from RNAi-EOF1 lead to the identification of additional proteins that could be regulated in EOF1 deficient eggshells. Our eggshell proteomics identified additional structural proteins and enzymes, suggesting mosquito extracellular eggshells are composed of a complex mixture of proteins. These function together to perform cross-linking, melanization, and sclerotization processes to create a protective shell capable of protecting the embryo and larva from environmental hazards for extended periods of time. Understanding reproductive processes of insect vectors of human disease could lead to the development of selective and safe small molecular inhibitors that may act to reduce the rate of disease transmission. These data provide new insights into mosquito ovarian maturation and eggshell synthesis that could lead to key advances in the field of vector control.

## Acknowledgments

We thank Dr. Peti Wolfgang at the Chemistry and Biochemistry Department (University of Arizona) for valuable advice on protein extraction methods from mosquito eggshells. We also thank Drs. Cynthia David and Krishna Parsawar for running eggshell proteomics at the Analytical and Biological Mass Spectrometry Core Facility in the Bio5 Institute, University of Arizona. Proteomics studies were funded internally through a Core Facilities Pilot Program provided by Research, Innovation & Impact at the University of Arizona (to J.I.).

## Materials and Methods

### Mosquitoes

*Aedes aegypti* (Rockefeller strain) were maintained on 10% sucrose and reared at 28°C, 80% relative humidity, and a 16 h light and 8 h dark cycle. Larvae in water were fed on a diet containing dry dog food, fish flakes, and liver powder (10:10:1 wt. ratio). Adult female mosquitoes were allowed to feed on blood supplemented with fresh adenosine triphosphate (5.0 mM final concentration) through stretched parafilm on an artificial glass feeder. Only fully engorged females identified under a light microscope were used. Human blood was donated by the American Red Cross (Tucson, AZ).

### RNAi screening of eggshell genes in *Aedes aegypti*

Target eggshell proteins were chosen from a proteomic analysis (Marinotti et al., 2014). RNA interference (RNAi) was performed to silence mRNA encoding mosquito eggshell genes (*SI Appendix*, Table S1). Each gene-specific forward and reverse oligonucleotide primer was designed using a NetPrimer web-based primer analysis tool. T7 RNA polymerase promoter sequence, TAATACGACTCACTATAGGGAGA, was placed to the 5′ end of each primer (*SI Appendix*, Table S2). All primers were obtained from Eurofins Genomics (Louisville, KY). PCR was performed using Taq 2X Master Mix (NEB, Ipswich, MA) with cDNA derived from mosquito whole body as a template, and the amplified PCR products were ligated into the pGEM-T easy vector (Promega Madison, WI) for DNA sequence verification. dsRNA was synthesized by *in vitro* transcription using the PCR products as templates, NTP nucleotides, and T7 RNA polymerase from HiScribe™ T7 Quick High Yield RNA Synthesis Kit (NEB). Cold-anesthetized female mosquitoes were microinjected with 2.0 µg gene-specific dsRNA using a Nanoject II microinjector (Drummond Scientific Company, Broomall, PA). Mosquitoes were maintained on 10% sucrose during the experiments. RNAi-mosquitoes were allowed to mate and feed a blood meal four days after dsRNA microinjection. Fully engorged females were individually placed into oviposition containers. Eggs on oviposition paper were photographed two days post oviposition under a light microscope (Nikon, SMZ-10A). Phenotypes including fecundity and egg melanization were recorded. To determine egg viability, eggs on oviposition papers remained wet for three days before drying at 28°C, and seven day-old eggs on oviposition paper were submerged in water, vacuumed using a Speed Vac for 10 minutes, and allowed to hatch for 2 days.

### Pattern of eggshell gene expression by quantitative real-time PCR (qPCR)

We determined expression patterns of four eggshell proteins, Nasrat, Closca, Polehole, and Nudel. Samples were obtained from larvae, pupae and male adults, as well as in five tissues including thorax, fat body, midgut, ovaries, and Malpighian tubules from sugar-fed at 3 day post eclosion and blood-fed female mosquitoes at 24 and 48 h PBM. Fourth instar larvae and pupae of mixed age and sexes were thoroughly washed with dH_2_O before placing them in TRIzol reagent (Invitrogen, Carlsbad, CA). Tissues were dissected in 1X PBS under a light microscope. Total RNA was extracted according to the manufacturer’s instructions. First strand cDNA was synthesized from DNaseI-treated total RNA using an oligo-(dT)_20_ primer and reverse transcriptase. qPCR was carried out with the corresponding cDNA, gene-specific primers (*SI Appendix*, Table S4), PerfeCTa SYBR Green FastMix, and ROX (Quanta BioSciences, Gaithersburg, MD) on the 7300 Real-Time PCR System (Applied Biosystems). Relative expression for Nasrat, Closca, Polehole, and Nudel was calculated as 2^−ΔΔCt^ and normalized to ribosomal protein S7 transcript levels in the same cDNA samples. Data were collected from three different biological cohorts of 10 individual mosquitoes.

### Measuring knockdown efficiency of eggshell genes at the transcription level

A level of RNAi-mediated transient knockdown was verified by qPCR using gene-specific primers (*SI Appendix*, Table S4). cDNA were synthesized from DNaseI-treated total RNA that were derived from ovaries of individual dsRNA-injected mosquitoes at 36 hours PBM. Relative expression for Nasrat, Closca, Polehole, and Nudel was calculated as 2^−ΔΔCt^ and normalized to ribosomal protein S7 transcript levels in the same cDNA samples. The knockdown efficiency was compared using Fluc dsRNA-injected mosquitoes as a control. Data were collected from 12 individual mosquitoes.

### *In vitro* follicle melanization assay

An *in vitro* follicle melanization assay was performed using mature primary follicles isolated from ovaries of RNAi mosquitoes (RNAi-Fluc, -Nasrat, -Closca, -Polehole, and -Nudel) at 96 hours PBM. The timing of dsRNA microinjection and blood feeding schedule are shown in Fig. 1. The follicles were placed on an oviposition paper wetted with distilled water and photographed 5, 90 and 150 min after dissection. Follicles from five individual mosquitoes were used. Dissected matured primary follicles were also soaked *in vitro* with protease inhibitors (PI, 1X concentration in water; cOmplete™ Mini Protease Inhibitor Cocktail, Sigma). Ovaries from female mosquitoes at 96 hours PBM were dissected and placed in water containing PI at different time points after ovary dissection. Individual follicles were then separated, transferred onto oviposition paper, continuously soaked with water containing the PI, and monitored for eggshell melanization. The control follicles were soaked in water without PI. Follicles were also treated with phenylmethylsulfonyl fluoride (PMSF) at a 10 mM final concentration.

### Rhodamine B mosquito follicle permeability assay

The advantage to using this type of assay is that it can quickly assess whether follicles within the ovaries may contain defective eggshell prior to oviposition. Ovaries of RNAi-mosquitoes were dissected at 96 hour PBM, and the individual primary mature follicles were separated from the ovaries and transferred to glass scintillation vials containing Rhodamine B (Sigma) at a 1.0 mM final concentration in water. The follicles were stained with Rhodamine B for 10 min on a rocking shaker and thoroughly rinsed with H_2_O. The follicles were photographed under a light microscope with a Coolpix 4300 (Nikon).

### Ultrastructural study of eggshell by SEM

The ovaries were dissected from mosquitoes that had undergone microinjection with different dsRNAs at 96 hours PBM. Mature primary follicles were separated from the ovaries in 1X PBS under a light microscope. The follicles were washed several times with 1X PBS and fixed with 2.5% EM-grade glutaraldehyde (EMS, Electron Microscopy Sciences, Hatfield, PA) in 0.1 M PIPES (pH 7.2) overnight at 4°C prior to being washed thoroughly with 0.1 M PIPES. The follicles were then post-fixed in 1% osmium tetroxide (EMS) in 0.1 M PIPES for 1 hour in the dark, and subsequently washed thoroughly in deionized water. Next, the follicles were dehydrated with ethanol (ETOH) in water through a graded series for 10 min each in 10, 30, 50, 70, 90% ETOH and three times for 30 min each in 100% ETOH at room temperature. The dehydrated follicles were then stored at 4°C for four days. The samples were dried with hexamethyldisilazane (HMDS, EMS) in ETOH through a graded series for 20 min each in 25, 50, 75, and 100% HMDS at room temperature. Finally, the follicle samples were air-dried under a fume hood overnight at room temperature prior to SEM analysis. The dried samples were metallized with gold using Hummer 6.2 Sputter System (Anatech USA, Union City, CA). An Inspect-S scanning electron microscope (FEI, Hillsboro, OR) was used to compare the ultrastructural surface features of the follicles isolated from various RNAi-mosquitoes at the W.M. Keck Center for Nano-Scale Imaging, University of Arizona.

### Mosquito eggshell protein extract preparation

Mosquitoes were microinjected with dsRNA against Fluc control or EOF1 one day after adult eclosion and allowed to feed on blood for four days after the injection, as shown in Fig. 1. Ovaries were dissected from the dsRNA injected mosquitoes four days post blood meal, and mature primary follicles were separated from dissected ovaries and thoroughly washed in 1X PBS in order to completely remove shed follicular epithelial cells, secondary follicles, the germarium, and other ovarian cells. The primary follicles were homogenized using a dounce homogenizer with pestle B, and the extracellular eggshells were filtered through a mesh strainer (40 μm) and washed thoroughly with distilled water to remove oocyte cytosolic and membrane contents. Next, the enriched eggshells were homogenized in 6.0 M guanidine hydrochloride using a dounce homogenizer with pestle A and incubated at 37°C overnight. The eggshell proteins were subsequently precipitated with 100% ethanol overnight at – 20°C and pelleted by a centrifugation (16,000 g for 15 min at 4°C). Finally, the eggshell proteins were resuspended with a protein sample buffer, quantified, and denatured in boiling water.

### Protein sample preparation for mass spectrometry

Protein sample preparation for mass spectrometry was carried out at the Analytical and Biological Mass Spectrometry Facility, University of Arizona. Eggshell protein samples were separated about 1.0 cm into resolving gel on precast SDS-12% PAGE gels (Bio Rad). The gels were fixed with 50% methanol and 10% glacial acetic acid for 30 min and stained with GelCode Blue Stain Reagent (Thermo Scientific) overnight. Excised gel bands were destained with 50/50 methanol/50 mM ammonium bicarbonate. Disulfide bonds were reduced with dithiothreitol (25 mM) for 45 min at 60°C and free cysteines were alkylated with iodoacetamide (55 mM) at room temperature for 30 min. After extensive washing with 50 mM ammonium bicarbonate, the bands were cut into small pieces and incubated with trypsin/lysC (Promega) digestion solution in 50 mM ammonium bicarbonate containing 0.1% ProteaseMAX (Promega) overnight at 37°C. Peptides were extracted from the gels with 45% ACN/5% IPA/0.2% formic acid by sonication for 10 min. The extraction was repeated once; supernatants were pooled and dried down in a speed vac.

### LC–MS/MS and protein identification

LC-MS/MS analysis was done on a Q Exactive Plus mass spectrometer (Thermo Fisher Scientific, San Jose, CA) equipped with an EASY-Spray nanoESI source. Peptides were eluted from an Acclaim Pepmap 100 trap column (75 micron ID × 2 cm, Thermo Scientific) onto an Acclaim PepMap RSLC analytical column (75 micron ID × 25 cm, Thermo Scientific) using a 3-20% gradient of solvent B (acetonitrile, 0.1% formic acid) over 120 min, 20-50% solvent B over 15 min, 50-95% of solvent B over 10 min, a hold of solvent 95% B for 10 min, and finally a return to 3% solvent B for 10 min. Solvent A consisted of water and 0.1% formic acid. Flow rates were 300 nL/min using a Dionex Ultimate 3000 RSLCnano System (Thermo Scientific). Data dependent scanning was performed by the Xcalibur v 4.0.27.19 software (Andon et al., 2002) using a survey scan at 70,000 resolution scanning mass/charge (m/z) 350-1600 at an automatic gain control (AGC) target of 1e6 and a maximum injection time (IT) of 65 msec, followed by higher-energy collisional dissociation (HCD) tandem mass spectrometry (MS/MS) at 27nce (normalized collision energy). Of the 11 most intense ions at a resolution of 17,500, an isolation width of 1.5 m/z, an AGC of 5e4 and a maximum IT of 65 msec were observed. Dynamic exclusion was set to place any selected m/z on an exclusion list for 30 seconds after a single MS/MS. Ions of charge state +1, 7, 8, >8, unassigned, and isotopes were excluded from MS/MS. MS and MS/MS data were searched against the NCBI *Aedes aegypti* protein database to which additional common contaminant proteins (e.g. trypsin, keratins; obtained at ftp://ftp.thegpm.org/fasta/cRAP) was appended using Thermo Proteome Discoverer v 2.2.0388 (Thermo Fisher Scientific). MS/MS spectra matches considered fully tryptic peptides with up to two missed cleavage sites. Variable modifications considered were methionine oxidation (15.995 Da), and cysteine carbamidomethylation (57.021 Da). Proteins were identified at 95% confidence with XCorr score cut-offs (Qian et al., 2005) as determined by a reversed database search. The protein and peptide identification results were further analyzed with Scaffold Q+S v 4.8.7 (Proteome Software Inc., Portland OR), a program that relies on various search engine results (i.e.: Sequest, X!Tandem, MASCOT) and which uses Bayesian statistics to reliably identify more spectra (Keller et al., 2002). Protein identifications were accepted that passed a minimum of two peptides identified at 0.1% peptide False Discovery Rate and 95% protein confidence by the Protein Profit algorithm within Scaffold. Only proteins with at least six matching peptide hits were included as eggshell proteins and subjected to protein BLAST searches against the NCBI non-redundant protein database to determine their putative protein functions.

### Statistical analysis

Statistical analyses were performed using GraphPad Prism Software (version 5.0; GraphPad, La Jolla, CA). Statistical significance for fecundity, melanization, viability, and RNAi knockdown efficiency was analyzed using an unpaired Student’s t-test. *P* values of ≤ 0.05 were considered significantly different. All experiments were performed using at least three independent biological cohorts except the proteomic studies.

## Supporting information (SI)

**Figure S1. Validation of the RNAi-mediated knockdown efficiency by quantitative real-time PCR (qPCR)**. Relative abundance of mRNA levels for Nasrat, Closca, Polehole, and Nudel was analyzed in dissected mosquito ovaries at 36 h PBM. Mosquitoes were microinjected with each dsRNA at four days prior to blood feeding, as shown in Fig. 1. dsRNA-Fluc-injected mosquitoes were used as controls. A single mosquito analysis was performed to isolate total RNA, synthesize cDNA, and monitor silencing efficiency by qPCR. mRNA levels were normalized according to transcript levels of ribosomal S7 protein. Data are presented as MEAN ± SEM of 12 individual mosquitoes. *** *P* < 0.001 compared to RNAi-Fluc. Vectorbase ID: Nasrat (AAEL008829), Closca (AAEL000961), Polehole (AAEL022628), and Nudel (AAEL016971). qPCR Primers used are shown in *SI Appendix*, Table S4.

**Figure S2. Validation of the RNAi-mediated knockdown efficiency by quantitative real-time PCR (qPCR)**. Relative abundance of mRNA levels for DCE2, DCE4, DCE5, and CATL3 was analyzed in dissected mosquito ovaries at 36 h PBM. Mosquitoes were microinjected with each dsRNA at four days prior to blood feeding, as shown in Fig. 1. dsRNA-Fluc-injected mosquitoes were used as controls. A single mosquito analysis was performed to isolate total RNA, synthesize cDNA, and monitor silencing efficiency by qPCR. mRNA levels were normalized according to transcript levels of ribosomal S7 protein. Data are presented as MEAN ± SEM of 12 individual mosquitoes. *** *P* < 0.001 compared to RNAi-Fluc. Vectorbase ID: DCE2 (AAEL006830), DCE4 (AAEL007096), DCE5 (AAEL010848), and CATL3 (AAEL002196). qPCR Primers used are shown in *SI Appendix*, Table S4.

**Video 1. A time-lapse video on a melanization process of eggs deposited by female *Aedes aegypti* mosquitoes that were microinjected with dsRNAi against Fluc control**. An oviposition paper was wetted with water periodically to prevent dryness. The video was taken and edited by Sony DCR-SR68 and Adobe Premiere Pro CS5, respectively.

**Video 2. Visualization of *Aedes aegypti* mosquito eggshell melanization in response to RNAi-Nudel by time-lapse video microscopy**. Eggs from female mosquitoes that were microinjected with dsRNA-Nudel were video recorded. An oviposition paper was wetted with water periodically to prevent dryness. The video was taken and edited by Sony DCR-SR68 and Adobe Premiere Pro CS5, respectively.

