## Supplementary material for "Characterization of essential eggshell proteins from *Aedes aegypti* mosquitoes": SI Appendix

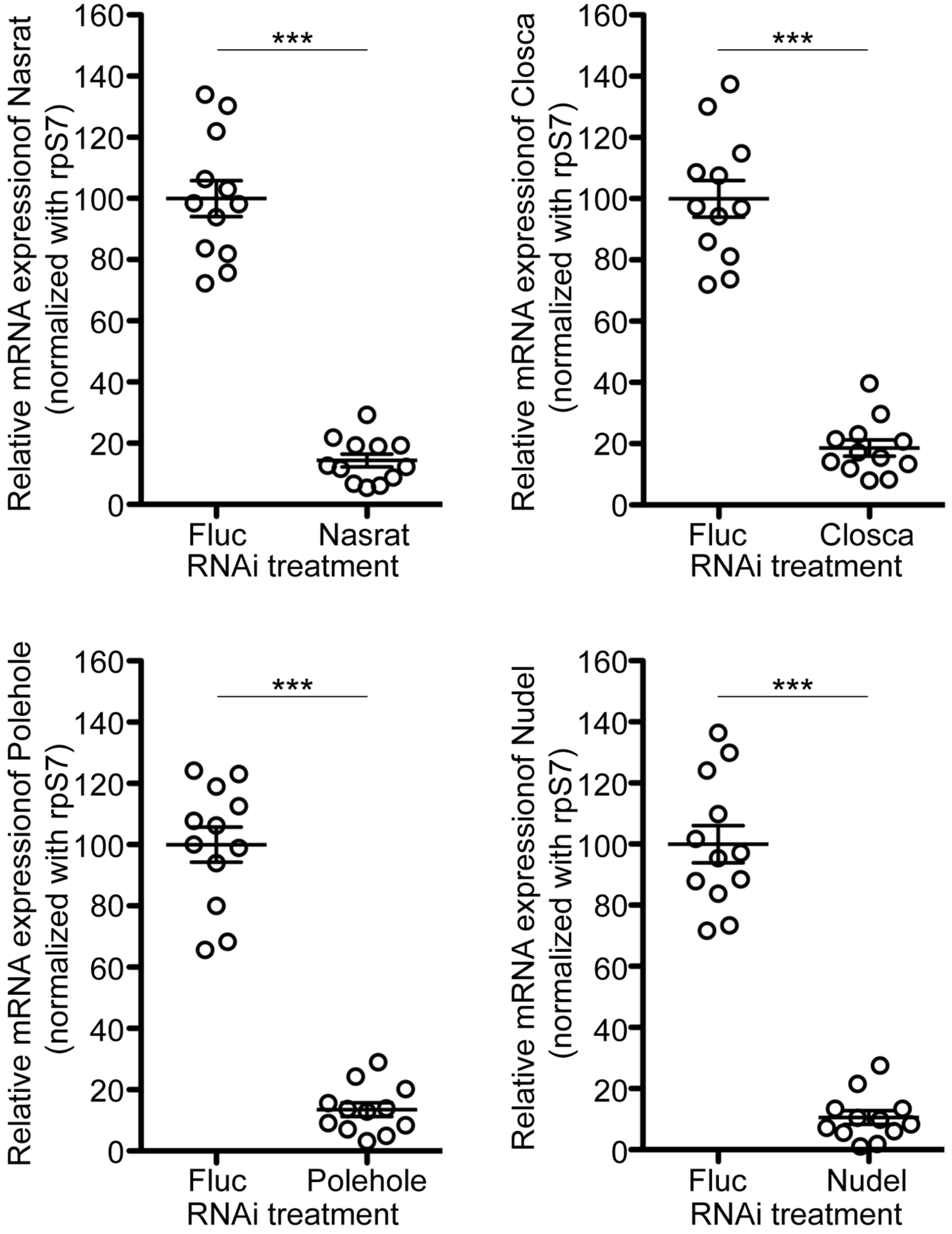


**Figure S1. Validation of the RNAi-mediated knockdown efficiency by quantitative real-time PCR (qPCR).** Relative abundance of mRNA levels for Nasrat, Closca, Polehole, and Nudel was analyzed in dissected mosquito ovaries at 36 h PBM. Mosquitoes were microinjected with each dsRNA at four days prior to blood feeding, as shown in Fig. 1. dsRNA-Fluc-injected mosquitoes were used as controls. A single mosquito analysis was performed to isolate total RNA, synthesize cDNA, and monitor silencing efficiency by qPCR. mRNA levels were normalized according to transcript levels of ribosomal S7 protein. Data are presented as MEAN ± SEM of 12 individual mosquitoes. *** $P < 0.001$ compared to RNAi-Fluc. Vectorbase ID: Nasrat (AAEL008829), Closca (AAEL000961), Polehole (AAEL022628), and Nudel (AAEL016971). qPCR Primers used are shown in *SI Appendix*, Table S4.

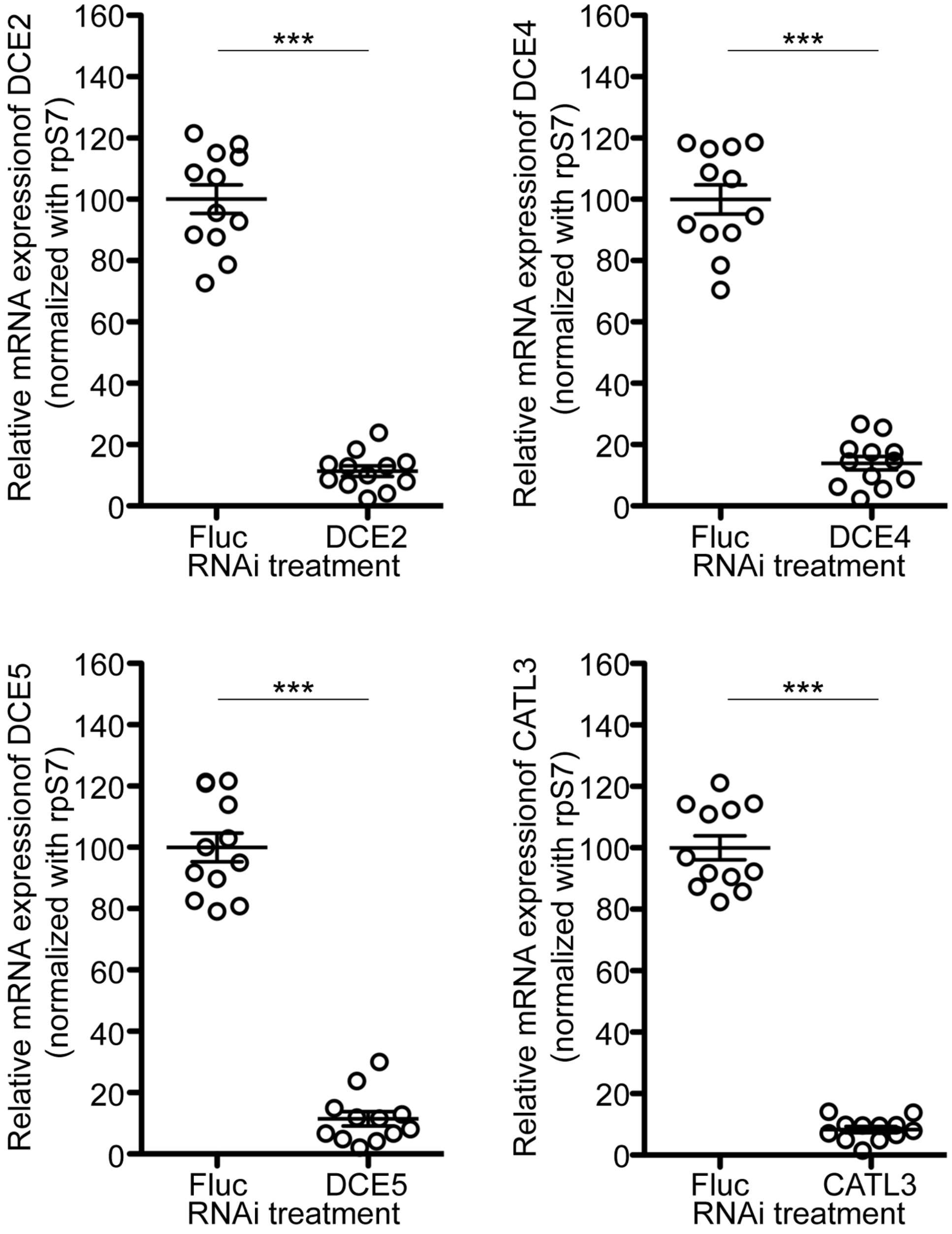


**Figure S2. Validation of the RNAi-mediated knockdown efficiency by quantitative real-time PCR (qPCR).** Relative abundance of mRNA levels for DCE2, DCE4, DCE5, and CATL3 was analyzed in dissected mosquito ovaries at 36 h PBM. Mosquitoes were microinjected with each dsRNA at four days prior to blood feeding, as shown in Fig. 1. dsRNA-Fluc-injected mosquitoes were used as controls. A single mosquito analysis was performed to isolate total RNA, synthesize cDNA, and monitor silencing efficiency by qPCR. mRNA levels were normalized according to transcript levels of ribosomal S7 protein. Data are presented as MEAN ± SEM of 12 individual mosquitoes. *** $P < 0.001$ compared to RNAi-Fluc. Vectorbase ID: DCE2 (AAEL006830), DCE4 (AAEL007096), DCE5 (AAEL010848), and CATL3 (AAEL002196). qPCR Primers used are shown in *SI Appendix*, Table S4.

Table S1. RNAi screening of *Aedes aegypti* eggshell proteins.

| Vectorbase ID | GenBank ID | Putative functions | *RNAi phenotypic effects* Eggs | Hatching |
|---|---|---|---|---|
| AAEL000361 | EAT48607 | Trypsin inhibitor-like/serpin | NO | NO |
| AAEL000363 | EAT48611 | Trypsin inhibitor-like/serpin | NO | NO |
| AAEL000375 | EAT48605 | Trypsin inhibitor-like/serpin | NO | NO |
| AAEL000507 | EAT48446 | Chorion peroxidase, CP4 | NO | NO |
| AAEL000961 | EAT47957 | Closca | YES | YES |
| AAEL002196 | EAT46597 | Cysteine proteinase L-like, CATL3 | YES | YES |
| AAEL002382 | EAT46452 | Unknown | NO | NO |
| AAEL003110 | EAT45649 | Chitinase domain | NO | NO |
| AAEL004202 | EAT44412 | Unknown | NO | NO |
| AAEL004386 | EAT44219 | Chorion peroxidase, CP1 | NO | NO |
| AAEL004390 | EAT44218 | Chorion peroxidase, CP2 | NO | NO |
| AAEL004401 | EAT44216 | Chorion peroxidase, CP7 | NO | NO |
| AAEL005098 | EAT43477 | Trypsin inhibitor-like/serpin | NO | NO |
| AAEL005648 | EAT42848 | Clip-domain serine protease | NO | NO |
| AAEL005861 | EAT42645 | Vacuolar sorting protein | NO | NO |
| AAEL006830 | EAT41553 | Dopachrome converting enzyme, DCE2 | YES | YES |
| AAEL006985 | EAT41324 | Dopachrome converting enzyme, DCE3 | NO | NO |
| AAEL007096 | EAT41240 | Dopachrome converting enzyme, DCE4 | YES | NO |
| AAEL007415 | EAT40867 | Laccase-like multicopper oxidases | NO | NO |
| AAEL007641 | EAT40646 | Transglutaminase | NO | NO |
| AAEL008829 | EAT39370 | Nasrat | YES | YES |
| AAEL009290 | EAT38853 | Unknown | NO | NO |
| AAEL009452 | EAT38674 | Unknown | NO | NO |
| AAEL009746 | EAT38349 | Unknown | NO | NO |
| AAEL010544 | EAT37465 | Unknown | NO | NO |
| AAEL010848 | EAT37145 | Dopachrome converting enzyme, DCE5 | YES | NO |
| AAEL011238 | EAT36701 | Trypsin inhibitor-like/serpin | NO | NO |
| AAEL012586 | EAT35235 | Unknown | NO | NO |
| AAEL013027 | EAT34764 | Vitelline membrane protein, 15a1 | NO | NO |
| AAEL013936 | EAT33799 | Trypsin inhibitor-like/serpin | NO | NO |
| AAEL014561 | EAT33176 | Vitelline membrane protein, 15a3 | NO | NO |
| AAEL015203 | EAT32616 | Unknown | NO | NO |
| AAEL017403 | EJY58008 | Vitelline membrane protein, 15a2 | NO | NO |
| AAEL017467 | EJY57339 | Chorion peroxidase, CP6 | NO | NO |

Table S2. Primers used for RNAi screening.

| Vectorbase ID | Gene-specific RNAi primers (5' - 3') | | Vectorbase ID | Gene-specific RNAi primers (5' - 3') | |
|---|---|---|---|---|---|
| AAEL000361 | Forward | GTAGCGATTGTTGTTCTAGCG | AAEL007415 | Forward | TCAGTGCCGACCAGCAAGT |
| | Reverse | GGCGTAAGTAACTTGCACACGG | | Reverse | CTGAGATTGTCTTGTTGGACTTC |
| AAEL000363 | Forward | CCCTGTGCCGACCCAAACGA | AAEL007641 | Forward | CGCTCGCAACGTGCATTGG |
| | Reverse | CGGTGTGGTCGTAGTAATACA | | Reverse | GGACACCACGCACGCTGCAA |
| AAEL000375 | Forward | ATGCAGCTTCCAATATGTGCTAT | AAEL008829 | Forward | GAGCCCATTCAGAACCTCCT |
| | Reverse | GCGGCTTCGGCGTAGGCTT | | Reverse | AGCGTAACTCCGTTGACGTA |
| AAEL000507 | Forward | TACAGCTCTGCGTGCATCTG | AAEL009290 | Forward | ATCGAGGGATTGATGGAAGG |
| | Reverse | CCACAATCGGTCTTCGTCAG | | Reverse | CCGTCCGAGTAGTGGATCGC |
| AAEL000961 | Forward | GGCAAGGGCTTCTACAACGT | AAEL009452 | Forward | GTCCTCCATCTCTTTGGTGA |
| | Reverse | CCGTTCAAAGTATGCTCCAC | | Reverse | TCACCAACCAGCTCTTCTCG |
| AAEL002196 | Forward | TGAAGAAACAACTGCTGTGG | AAEL009746 | Forward | CAGCCTACATCGTTGACCTA |
| | Reverse | CCATCTGCTGGGTACTGAC | | Reverse | TGTCGAAGCACAACCGATGGTT |
| AAEL002382 | Forward | GCCCATGTGAACTTCCCTTGCC | AAEL010544 | Forward | CAGCGGGATCAGAACCAGGAT |
| | Reverse | GATACCTCGCCCTGTTGAAC | | Reverse | CATAAGATCCGTCAGACCGTC |
| AAEL003110 | Forward | TCCAACCAGGAGTCGAGTGA | AAEL010848 | Forward | GCCCTCGACTCAGGCATTTG |
| | Reverse | CGCCAATTCCACCGAGTTG | | Reverse | TGCGTGCTCAAGCGACACTC |
| AAEL004202 | Forward | TCCTACGGCGAAGCTGGTTC | AAEL011238 | Forward | CAACCAGTTGATGGCAGGATAC |
| | Reverse | GACTCTCGTTTGTCTGCTTC | | Reverse | CCGTTGTGCTTCACATAACC |
| AAEL004386 | Forward | TGAGGGAACACAACCGACTA | AAEL012586 | Forward | GCCGACAGGGACCGATGATG |
| | Reverse | TAAACCTGTGCCAAGAGTGC | | Reverse | GCCGAAATGTTGATCTTGTGTAC |
| AAEL004390 | Forward | TGAGGGAACACAACCGACTA | AAEL013027 | Forward | TTCCCATCCAACTCAGTAACCAT |
| | Reverse | TAAACTCGCGCCAGAAGAGC | | Reverse | TTCCGCTGCATCTTCAAGAG |
| AAEL004401 | Forward | CCACACTGGTCTGACGACAT | AAEL013936 | Forward | TTAGCAATAGTTTCTCACTGCCA |
| | Reverse | CGCCTACGTAAAGATCGACGT | | Reverse | GCCTGTGGGCTTCGATTGG |
| AAEL005098 | Forward | ATGAAGTTGGCAATCATTTGTGT | AAEL014561 | Forward | CGGAAGGAATCCATCCAACTT |
| | Reverse | GCTCCAGGACAATCGCACAG | | Reverse | CAGTCCAATCGATGATCCGC |
| AAEL005648 | Forward | GCCAAAGCCGATAGCCATC | AAEL015203 | Forward | GTGTTGGTGCCGAAGAAGAG |
| | Reverse | CTAGGCATGTTGAGAGCACC | | Reverse | TAGCACTTCAACTCGGATGACTT |
| AAEL005861 | Forward | TGTGATGGCGATGACGACTG | AAEL017403 | Forward | CCAGCGTGGTACAACAGTAAATC |
| | Reverse | CTCATCACTTCCATCCTTGCA | | Reverse | CCGTTCCTTGGTCCTGGTTC |
| AAEL006830 | Forward | TGTGGAAATCGTCGGTGGT | AAEL017467 | Forward | TACAGCTCTGCGTGCATCTG |
| | Reverse | TGTAGGCGAAGGTGTCCTC | | Reverse | CCACAATCGGTCTTCGTCAG |
| AAEL006985 | Forward | CTCCCGGTTGGAATCGAAAG | AAEL012336 | Forward | AGCCCGTCCAAGAGGAAGTT |
| | Reverse | GTCCGTAGTCCAGTTCATTGG | (EOF1) | Reverse | CTCGGATGGTACTCACACAA |
| AAEL007096 | Forward | GCAAGAAGTGGCGACAAGAC | U47295 | Forward | AGCACTCTGATTGACAAATACGA |
| | Reverse | CGTCCACCCAGATAGGTGAA | (Luciferase) | Reverse | AGTTCACCGGCGTCATCGTC |

T7 promoter sequence (5' TAATACGACTCACTATAGGGAGA 3') was added in 5' of each RNAi primer.

Table S3. Reproductive phenotypes associated with RNAi in *Aedes aegypti.*

| RNAi | Fluc | Nasrat | Closca | Polehole | Nudel |
|---|---|---|---|---|---|
| *Fecundity* | | | | | |
| Number of mosquitoes examined | 24 | 26 | 25 | 27 | 24 |
| Total number of eggs oviposited | 2273 | 2128 | 2073 | 2359 | 1955 |
| Mean number of eggs oviposited | 94.7 | 81.8 | 82.9 | 87.4 | 81.5 |
| *Eggshell melanization* | | | | | |
| Number of eggs examined | 2273 | 2128 | 2073 | 2359 | 1955 |
| Incompletely tanned eggs oviposited | 23 | 1520 | 1749 | 1955 | 1938 |
| Incomplete eggshell melanization (%) | 1.01% | 79.89% | 84.37% | 82.87% | 99.13% |
| *Egg viability* | | | | | |
| Number of eggs examined | 666 | 2128 | 2073 | 2359 | 1955 |
| Number of eggs hatched | 626 | 231 | 184 | 226 | 5 |
| Egg viability (%) | 93.99% | 10.86% | 8.88% | 9.58% | 0.26% |

Egg phenotypes are shown in Figure 2.

dsRNA was microinjected 4 days prior to blood feeding as shown in Figure 1.

Table S4. Gene-specific primers used for RNAi and qPCR in *Aedes aegypti.*

| Genes | | Primer sequence (5' to 3') |
|---|---|---|
| *Gene-specific primers used for RNAi* | | |
| Polehole, AAEL022628 | Forward | TGTACGGAAGGCAGGATTC |
| | Reverse | GGGTAGAGTTTGGTCAGGTT |
| Nudel, AAEL016971 | Forward | GCAAGCTGATGAAGTTCCACA |
| | Reverse | CCACGTCTTGTTCTCGGTTTC |
| *Gene-specific primers used for qPCR* | | |
| Nasrat | Forward | CTGAACACCGATCAGACGAT |
| | Reverse | TGAGCGTATTCTTGCGTTCGTA |
| Closca | Forward | GCCACCGACGTGCTGTTGAA |
| | Reverse | TCCCGTAGTTTAGCGTAGTTC |
| Polehole | Forward | ATAGTTCATCAGTTTCGATGCC |
| | Reverse | AACATCACATCGAACAAGAGTGT |
| Nudel | Forward | CACTTCGAGAACCAACATAAGG |
| | Reverse | GGAAGGTGATGTGCGTTAG |
| DCE2 | Forward | GCTAACATTGCCATCGACATG |
| | Reverse | GCCAGGACTTGTTCTTCTCAA |
| DCE4 | Forward | TGTAGATTCCGCGACGTTCT |
| | Reverse | CGTAGGTGATCCGTAGCAAT |
| DCE5 | Forward | TGGAACACTGATCAACCGTACAA |
| | Reverse | GACAATGAGTACATCAGAGCATC |
| CATL3 | Forward | GCCCTCAATGGACAGATTATG |
| | Reverse | GATCCTCCAGCACATCCCTT |
| Ribsomal protein S7 | Forward | ACCGCCGTCTACGATGCCA |
| | Reverse | ATGGTGGTCTGCTGGTTCTT |

T7 promoter sequence (5' TAATACGACTCACTATAGGGAGA 3') was added in 5' of each RNAi primer.

Table S5. Reproductive phenotypes associated with RNAi in *Aedes aegypti*

| RNAi | Fluc | Nasrat | Closca | Polehole | Nudel |
|---|---|---|---|---|---|
| *Fecundity* | | | | | |
| Number of mosquitoes examined | 12 | 12 | 12 | 12 | 12 |
| Total number of eggs oviposited | 1076 | 1012 | 1022 | 1024 | 976 |
| Mean number of eggs oviposited | 89.7 | 84.3 | 85.2 | 85.3 | 81.3 |
| *Eggshell melanization* | | | | | |
| Number of eggs examined | 1076 | 1012 | 1022 | 1024 | 976 |
| Incompletely tanned eggs oviposited | 14 | 30 | 22 | 31 | 964 |
| Incomplete eggshell melanization (%) | 1.30% | 2.96% | 2.15% | 3.03% | 98.77% |
| *Egg viability* | | | | | |
| Number of eggs examined | 364 | 368 | 348 | 342 | 976 |
| Number of eggs hatched | 333 | 332 | 312 | 302 | 8 |
| Egg viability (%) | 91.48% | 90.22% | 89.66% | 88.30% | 0.82% |

Egg phenotypes are shown in Fig. 4.

dsRNA was microinjected immediately after blood feeding as shown in Fig. 4.

Table S6. Reproductive phenotypes associated with RNAi in two gonotrophic cycles.

| | First gonotrophic cycle | | Second gonotrophic cycle | |
|---|---|---|---|---|
| RNAi | Fluc | Nudel | Fluc | Nudel |
| *Fecundity* | | | | |
| Number of mosquitoes examined | 12 | 12 | 12 | 12 |
| Total number of eggs oviposited | 1007 | 902 | 807 | 731 |
| Mean number of eggs oviposited | 83.9 | 75.2 | 67.3 | 60.9 |
| *Eggshell melanization* | | | | |
| Number of eggs examined | 1007 | 902 | 807 | 731 |
| Incompletely tanned eggs oviposited | 11 | 890 | 22 | 21 |
| Incomplete eggshell melanization (%) | 1.09% | 98.67% | 2.73% | 2.87% |
| *Egg viability* | | | | |
| Number of eggs examined | 365 | 902 | 383 | 393 |
| Number of eggs hatched | 335 | 7 | 338 | 337 |
| Egg viability (%) | 91.78% | 0.78% | 88.25% | 85.75% |

Egg phenotypes are shown in Fig. 5.

dsRNA was microinjected immediately after blood feeding as shown in Fig. 5.

Table S7. An *in vitro* follicle melanization assay in *Aedes aegypti*.

| *RNAi treatment* | Fluc | Nasrat | Closca | Polehole | Nudel |
|---|---|---|---|---|---|
| Number of mosquitoes examined | 5 | 5 | 5 | 5 | 5 |
| Total number of follicles examined | 118 | 121 | 105 | 107 | 113 |
| Total number of follicles melanized | 114 | 6 | 13 | 15 | 4 |
| Follicle melanized (%) | 96.6% | 5.0% | 12.4% | 14.0% | 3.5% |

| *Protease inhibitor (PI) treatment on wildtype mosquitoes* | | PI added minutes after follicle dissection | | |
|---|---|---|---|---|
| | Untreated | 0 | 10 | 20 |
| Number of mosquitoes examined | 5 | 5 | 5 | 5 |
| Total number of follicles examined | 122 | 146 | 130 | 132 |
| Total number of follicles melanized | 118 | 3 | 125 | 126 |
| Follicle melanized (%) | 96.7% | 2.1% | 96.2% | 95.5% |

Follicle phenotypes are shown in Figure 6.
dsRNA was microinjected 4 days prior to blood feeding as shown in Figure 1.

Table S8. An *in vitro* follicle melanization assay in *Aedes aegypti*.

| *RNAi treatment* | Fluc | Nasrat | Closca | Polehole | Nudel |
|---|---|---|---|---|---|
| Number of mosquitoes examined | 5 | 5 | 5 | 5 | 5 |
| Total number of follicles examined | 127 | 133 | 123 | 123 | 117 |
| Total number of follicles stained | 3 | 126 | 119 | 118 | 113 |
| Follicles stained (%) | 2.4% | 94.7% | 96.7% | 95.9% | 96.6% |

Follicle stained phenotypes are shown in Figure 6.
dsRNA was microinjected 4 days prior to blood feeding as shown in Figure 1.

Table S9. Reproductive phenotypes associated with RNAi in *Aedes aegypti.*

| RNAi | Fluc | DCE2 | DCE4 | DCE5 | CATL3 |
|---|---|---|---|---|---|
| *Fecundity* | | | | | |
| Number of mosquitoes examined | 23 | 24 | 25 | 25 | 26 |
| Total number of eggs oviposited | 2022 | 2128 | 2144 | 2130 | 2195 |
| Mean number of eggs oviposited | 87.9 | 88.7 | 85.8 | 85.2 | 84.4 |
| *Eggshell melanization* | | | | | |
| Number of eggs examined | 2022 | 2128 | 2144 | 2130 | 2195 |
| Incompletely tanned eggs oviposited | 19 | 2036 | 27 | 36 | 952 |
| Incomplete eggshell melanization (%) | 0.94% | 95.68% | 1.26% | 1.69% | 43.37% |
| *Egg viability* | | | | | |
| Number of eggs examined | 605 | 751 | 685 | 714 | 767 |
| Number of eggs hatched | 557 | 45 | 623 | 630 | 33 |
| Egg viability (%) | 92.07% | 5.99% | 90.95% | 88.24% | 4.30% |

Egg phenotypes are shown in Figure 7.

dsRNA was microinjected 4 days prior to blood feeding as shown in Figure 1.

Table S10. Raw data on eggshell proteomic analyses in *Aedes aegypti*.

| | | | # peptide hits | |
|---|---|---|---|---|
| Vectorbase ID | GenBank ID | Putative functions | RNAi-Fluc | RNAi-EOF1 |
| AAEL010434 | AAA18221 | Vitellogenin | 244 | 323 |
| AAEL006126 | EAT42292 | Vitellogenin | 281 | 222 |
| AAEL006138 | XP_001657509 | Vitellogenin | 278 | 219 |
| AAEL006830 | XP_001658066 | Dopachrome converting enzyme | 180 | 230 |
| AAEL004390 | XP_001649029 | Chorion peroxidase | 153 | 170 |
| AAEL010872 | XP_001661124 | Odorant binding protein | 172 | 141 |
| AAEL026038 | XP_021692994 | Chorion peroxidase | 160 | 146 |
| AAEL000961 | XP_021708553 | Closca | 143 | 161 |
| AAEL013492 | EAT34242 | Phenoloxidase | 156 | 147 |
| AAEL008829 | XP_001659577 | Nasrat | 112 | 128 |
| AAEL007415 | EAT40867 | Multicopper oxidase | 87 | 106 |
| AAEL004386 | P82600 | Chorion peroxidase | 100 | 82 |
| AAEL011763 | XP_001661890 | Phenoloxidase | 93 | 78 |
| AAEL022628 | XP_021707815 | Polehole | 67 | 88 |
| AAEL003110 | EAT45649 | Chitinase | 86 | 61 |
| AAEL003511 | XP_001656921 | Odorant binding protein | 108 | 33 |
| AAEL009433 | XP_001649879 | Odorant binding protein | 95 | 42 |
| AAEL009599 | XP_001660278 | Odorant binding protein | 98 | 37 |
| AAEL022918 | XP_021710656 | Glucose dehydrogenase | 65 | 61 |
| AAEL006396 | EAT42032 | Odorant binding protein | 90 | 35 |
| AAEL002851 | ABH03477 | Tubulin beta | 56 | 63 |
| AAEL026563 | XP_021695347 | Odorant binding protein | 66 | 42 |
| AAEL005752 | XP_001651411 | Lysosomal alpha-mannosidase | 61 | 43 |
| AAEL006563 | P42660 | Vitellogenic carboxypeptidase | 42 | 55 |
| AAEL006393 | EAT42030 | Odorant binding protein | 74 | 23 |
| AAEL008797 | XP_001659527 | Titin | 61 | 33 |
| AAEL005325 | EAT43230 | Dopachrome converting enzyme | 53 | 40 |
| AAEL006642 | XP_001652144 | Tubulin alpha-1 | 43 | 47 |
| AAEL000796 | EAT48134 | Odorant binding protein | 55 | 31 |
| AAEL006398 | EAT42033 | Odorant binding protein | 57 | 28 |
| AAEL005198 | EAT43356 | Juvenile hormone esterase | 49 | 35 |
| AAEL016971 | EJY57924 | Nudel | 54 | 26 |
| AAEL009642 | XP_001653891 | Cathepsin B-like cysteine proteinase 3 | 57 | 23 |
| AAEL007096 | XP_001658111 | Dopachrome converting enzyme | 42 | 35 |
| AAEL013338 | XP_001663495 | Lethal(2)essential for life | 43 | 31 |
| AAEL017467 | EJY57339 | Chorion peroxidase | 48 | 25 |
| AAEL011758 | ABF18058 | Cyclophylin | 42 | 27 |
| AAEL011197 | AAY81972 | Actin | 34 | 31 |
| AAEL011116 | XP_001655111 | 14-3-3 protein epsilon | 32 | 27 |
| AAEL019403 | ABF18332 | Heat shock 70 | 40 | 14 |
| AAEL022697 | XP_021706511 | Odorant binding protein | 40 | 14 |
| AAEL007339 | XP_001652677 | Heat shock protein 67b2 | 32 | 22 |
| AAEL006328 | XP_021710623 | Chitinase | 38 | 14 |
| AAEL000889 | XP_001651634 | Carboxylic ester hydrolase | 34 | 18 |
| AAEL003315 | EAT45429 | Odorant binding protein | 30 | 21 |
| AAEL005733 | XP_021701099 | Myosin heavy chain | 12 | 38 |
| AAEL009895 | EAT38188 | Neprilysin | 36 | 14 |
| AAEL017096 | XP_011493435 | Elongation factor 1-alpha | 30 | 19 |
| AAEL000144 | EAT48853 | Chitinase | 39 | 10 |
| AAEL000377 | EAT48658 | Odorant binding protein | 30 | 18 |
| AAEL015116 | AAG02219 | Prophenoloxidase | 15 | 31 |
| AAEL012062 | XP_021693479 | Na+/K+ ATPase alpha subunit | 14 | 32 |
| AAEL013719 | EAT34018 | Odorant binding protein | 29 | 16 |
| AAEL015289 | EAT32570 | Uncharacterized protein | 18 | 25 |
| AAEL004434 | XP_001649141 | Transketolase-like protein 2 | 18 | 24 |

Continued on next page

Table S10. Raw data on eggshell proteomic analyses in *Aedes aegypti* (continued).

| Vectorbase ID | GenBank ID | Putative functions | # peptide hits RNAi-Fluc | # peptide hits RNAi-EOF1 |
|---|---|---|---|---|
| AAEL010848 | XP_021698010 | Dopachrome converting enzyme | 32 | 10 |
| AAEL013501 | EAT34239 | Prophenoloxidase | 20 | 22 |
| AAEL000507 | EAT48446 | Chorion peroxidase | 33 | 9 |
| AAEL017501 | EJY57337 | NA-vitelline membrane | 17 | 24 |
| AAEL011708 | EAT36186 | Heat shock protein 83 | 28 | 13 |
| AAEL014876 | EAT32887 | Odorant binding protein | 28 | 12 |
| AAEL003393 | EAT45330 | ATP synthase beta subunit | 26 | 13 |
| AAEL022484 | XP_021706407 | Odorant binding protein | 25 | 14 |
| AAEL000318 | EAT48660 | Odorant binding protein | 20 | 19 |
| AAEL000833 | XP_001651310 | Odorant binding protein | 23 | 16 |
| AAEL015312 | EAT32556 | Cysteine proteinase-1 | 22 | 16 |
| AAEL004172 | EAT44445 | Alpha-Tubulin | 16 | 22 |
| AAEL014431 | EAT33288 | Odorant binding protein | 22 | 15 |
| AAEL004028 | XP_001648297 | Glucose dehydrogenase | 25 | 12 |
| AAEL001174 | EAT47749 | Odorant binding protein | 28 | 8 |
| AAEL009955 | XP_021707150 | Apolipophorins | 6 | 29 |
| AAEL000344 | XP_001655549 | Odorant binding protein | 20 | 15 |
| AAEL009038 | XP_021697410 | Serine protease/prolylcarboxypeptidase | 23 | 12 |
| AAEL007599 | EAT40702 | Cathepsin B | 27 | 7 |
| AAEL002023 | EAT46826 | Chitinase | 19 | 14 |
| AAEL014430 | EAT33287 | Odorant binding protein | 20 | 11 |
| AAEL006820 | XP_001658058 | Lipid storage droplets-binding protein 2 | 24 | 7 |
| AAEL011300 | EAT36638 | Uncharacterized protein | 17 | 14 |
| AAEL004500 | AAK01430 | Eukaryotic translation elongation | 17 | 13 |
| AAEL006336 | EAT42103 | Chitinase | 28 | 2 |
| AAEL008787 | O16109 | V-ATPase subunit A | 15 | 15 |
| AAEL014222 | EAT33503 | Vitellogenin receptor | 17 | 12 |
| AAEL024387 | XP_021712253 | Serine protease | 19 | 10 |
| AAEL018219 | XP_021705132 | BM-specific heparan sulfate proteoglycan | 11 | 17 |
| AAEL005766 | XP_001651423 | Fructose-bisphosphate aldolase | 21 | 7 |
| AAEL001593 | EAT47332 | GAPDH 1 | 14 | 13 |
| AAEL006271 | EAT42157 | Superoxide dismutase 3 | 15 | 11 |
| AAEL005756 | EAT42731 | Uncharacterized protein | 20 | 6 |
| AAEL012175 | XP_001655906 | ATP synthase alpha subunit | 16 | 9 |
| AAEL001061 | XP_011493351 | Glutathione S transferase | 14 | 11 |
| AAEL005759 | EAT42734 | Uncharacterized protein | 21 | 4 |
| AAEL013359 | XP_001656676 | DEAD box ATP-dependent RNA helicase | 11 | 14 |
| AAEL000496 | XP_001656955 | Chorion peroxidase | 15 | 10 |
| AAEL006322 | EAT42110 | Odorant binding protein | 11 | 14 |
| AAEL001189 | EAT47748 | Uncharacterized protein | 13 | 11 |
| AAEL010912 | EAT37053 | DPPIV-SP, Omega | 12 | 12 |
| AAEL013279 | XP_001663442 | Peptidyl-prolyl cis-trans isomerase | 14 | 9 |
| AAEL017502 | EJY57472 | Uncharacterized protein | 20 | 3 |
| AAEL010874 | EAT37093 | Odorant binding protein | 20 | 2 |
| AAEL007097 | XP_001658126 | 4-nitrophenylphosphatase | 15 | 7 |
| AAEL022038 | XP_021695232 | Uncharacterized protein | 15 | 6 |
| AAEL000703 | XP_001650265 | Glycogen phosphorylase | 17 | 4 |
| AAEL003513 | EAT45173 | Uncharacterized protein | 16 | 4 |
| AAEL000179 | XP_001658903 | Ubiquitin-conjugating enzyme E2 | 11 | 9 |
| AAEL027038 | XP_021710733 | Uncharacterized protein | 11 | 9 |
| AAEL008799 | XP_001659528 | Uncharacterized protein | 15 | 5 |
| AAEL010097 | XP_001654240 | Maternal protein exuperantia | 12 | 7 |
| AAEL023983 | ABF18051 | 40S ribosomal protein S4 | 9 | 10 |
| AAEL012054 | EAT35810 | Uncharacterized protein | 11 | 8 |
| AAEL013353 | XP_001656670 | Chicadee/profilin | 7 | 12 |

Continued on next page

Table S10. Raw data on eggshell proteomic analyses in *Aedes aegypti* (continued).

| Vectorbase ID | GenBank ID | Putative functions | # peptide hits RNAi-Fluc | # peptide hits RNAi-EOF1 |
|---|---|---|---|---|
| AAEL002160 | XP_001654823 | GTP-binding protein | 11 | 8 |
| AAEL003094 | EAT45648 | Chitinase | 15 | 4 |
| AAEL026426 | XP_021695306 | Vitelline membrane protein-like | 10 | 8 |
| AAEL009207 | EAT38962 | Uncharacterized protein | 13 | 5 |
| AAEL02734 | XP_021694355 | Uncharacterized protein | 11 | 7 |
| AAEL012960 | XP_001663145 | Pendulin (NLS-receptor) | 14 | 4 |
| AAEL007432 | EAT40884 | Serine protease | 18 | 0 |
| AAEL008862 | XP_001653480 | Metalloprotease | 14 | 3 |
| AAEL001863 | AAT36732 | Zinc carboxypeptidase | 12 | 5 |
| AAEL001153 | EAT47747 | Uncharacterized protein | 14 | 3 |
| AAEL005216 | EAT43331 | Uncharacterized protein | 14 | 3 |
| AAEL001588 | XP_001653563 | Glutamate carboxypeptidase | 12 | 5 |
| AAEL011808 | EAT36073 | Gglucose dehydrogenase | 11 | 6 |
| AAEL008640 | EAT39566 | Odorant binding protein | 13 | 4 |
| AAEL016984 | XP_011493026 | Glyceraldehyde-3-phosphate dehydrogenase | 11 | 5 |
| AAEL004388 | EAT44220 | Chorion peroxidase | 0 | 16 |
| AAEL004516 | XP_001649345 | Odorant binding protein | 9 | 7 |
| AAEL015288 | EAT32569 | Uncharacterized protein | 8 | 8 |
| AAEL006885 | XP_001652301 | 14-3-3 zeta | 9 | 7 |
| AAEL019604 | XP_021698211 | Uncharacterized protein | 10 | 5 |
| AAEL010585 | XP_001654680 | TER94 | 15 | 0 |
| AAEL010821 | XP_001655016 | 60S ribosomal protein LP0 | 8 | 7 |
| AAEL012996 | XP_021709703 | Rho guanine dissociation factor | 10 | 5 |
| AAEL008500 | XP_001659287 | DEAD box ATP-dependent RNA helicase | 11 | 3 |
| AAEL001965 | ABF18180 | Chitinase/imaginal disc growth factor | 5 | 9 |
| AAEL005901 | XP_001663344 | 40S ribosomal protein S3a | 8 | 6 |
| AAEL020238 | Q16ZR8 | 40S ribosomal protein SA | 10 | 4 |
| AAEL001845 | EAT47019 | Sepiapterin reductase | 8 | 6 |
| AAEL003525 | EAT45172 | Odorant binding protein | 14 | 0 |
| AAEL005422 | XP_001650875 | Pyrroline-5-carboxylate dehydrogenase | 11 | 3 |
| AAEL004984 | XP_001650136 | Cullin-associated NEDD8-dissociated protein | 14 | 0 |
| AAEL000827 | EAT48139 | Odorant binding protein | 10 | 4 |
| NA | CAF02084 | Odorant binding protein | 12 | 2 |
| AAEL003872 | XP_001664282 | Translationally-controlled tumor protein | 8 | 6 |
| AAEL001179 | EAT47746 | Odorant binding protein | 10 | 4 |
| AAEL009994 | XP_001660544 | 60S ribosomal protein L4 | 9 | 4 |
| AAEL012897 | XP_001663037 | Aconitase, mitochondrial | 10 | 3 |
| AAEL009496 | XP_001660169 | 40S ribosomal protein S7 | 6 | 7 |
| AAEL014548 | EAT33191 | Thioredoxin peroxidase | 6 | 7 |
| AAEL019408 | AAL37254 | 2-Cys thioredoxin peroxidase | 9 | 4 |
| AAEL028058 | XP_021695267 | Odorant binding protein | 9 | 4 |
| AAEL012035 | XP_001655825 | Vacuolar ATP synthase subunit e | 7 | 6 |
| AAEL014274 | XP_001648333 | Uncharacterized protein | 7 | 6 |
| AAEL000821 | EAT48127 | Odorant binding protein | 11 | 2 |
| AAEL017349 | XP_011493320 | HSP70 | 8 | 4 |
| AAEL019579 | XP_021704200 | Furin-like protease 2 | 7 | 5 |
| AAEL001432 | EAT47483 | Protein disulfide isomerase | 9 | 3 |
| AAEL009097 | EAT39077 | Cathepsin | 7 | 5 |
| AAEL004978 | XP_001650127 | DEAD box ATP-dependent RNA helicase | 9 | 3 |
| AAEL003820 | EAT44826 | Histone H2A-like | 6 | 6 |
| AAEL000758 | EAT48168 | Ubiquitin activating enzyme 1 | 11 | 1 |
| AAEL017451 | XP_011493087 | Angiotensin converting enzyme | 9 | 3 |
| AAEL009077 | EAT39089 | Alkaline phosphatase | 9 | 3 |
| AAEL008280 | EAT39971 | Uncharacterized protein | 5 | 6 |
| AAEL010168 | XP_001654299 | 40S ribosomal protein S2 | 6 | 5 |

Continued on next page

Table S10. Raw data on eggshell proteomic analyses in *Aedes aegypti* (continued).

| Vectorbase ID | GenBank ID | Putative functions | # peptide hits | |
|---|---|---|---|---|
| | | | RNAi-Fluc | RNAi-EOF1 |
| AAEL001495 | EAT47408 | Uncharacterized protein | 2 | 9 |
| AAEL019935 | XP_021709761 | Midline fasciclin | 7 | 4 |
| AAEL000837 | EAT48137 | Odorant binding protein | 7 | 4 |
| AAEL008481 | XP_001659268 | 60S ribosomal protein L18 | 6 | 5 |
| AAEL000846 | XP_001651309 | Odorant binding protein | 13 | 8 |
| AAEL011870 | XP_001662014 | Trailer hitch,protein LSM14 homolog B | 9 | 2 |
| AAEL010919 | EAT37047 | Prophenoloxidase | 5 | 6 |
| AAEL007003 | EAT41362 | Odorant binding protein | 7 | 4 |
| AAEL008381 | EAT39841 | Peptide transporter | 5 | 6 |
| AAEL004532 | EAT44083 | Glyoxylate reductase | 6 | 5 |
| AAEL005798 | XP_001651458 | ATP synthase subunit beta vacuolar | 6 | 5 |
| AAEL002978 | EAT45789 | Aminopeptidase | 8 | 3 |
| AAEL012904 | XP_001663045 | Rab gdp-dissociation inhibitor | 8 | 3 |
| AAEL014719 | EAT33023 | Uncharacterized protein | 7 | 4 |
| AAEL007962 | XP_001658750 | Glutathione transferase | 7 | 4 |
| AAEL004755 | EAT43826 | Enoyl-CoA delta isomerase | 10 | 1 |
| AAEL014238 | AAC31639 | DOPA decarboxylase | 9 | 1 |
| AAEL011288 | ABF18271 | Eukaryotic translation elongation factor 1-$\gamma$ | 8 | 2 |
| AAEL009287 | XP_001659895 | Ran, GTP-binding nuclear protein | 6 | 4 |
| AAEL006389 | ABE72972 | Cathepsin L | 8 | 2 |
| AAEL007915 | Q170J7 | Moesin | 7 | 3 |
| AAEL004856 | XP_001649919 | Odorant binding protein | 8 | 2 |
| AAEL001487 | EAT47390 | Odorant binding protein | 7 | 3 |
| AAEL006977 | XP_001652452 | Ser/thr protein phosphatase 2a reg. subunit a | 8 | 2 |
| AAEL009882 | XP_001654079 | Retinoblastoma-binding protein 4 | 6 | 4 |
| AAEL011764 | EAT36127 | Phenoloxidase | 2 | 8 |
| AAEL003404 | XP_001656805 | Uncharacterized protein | 0 | 10 |
| AAEL022214 | XP_021707151 | $\gamma$-interferon inducible lys. thiol reductase | 7 | 3 |
| AAEL007236 | ABF18366 | Uncharacterized protein | 6 | 4 |
| AAEL000987 | XP_001657711 | 60S ribosomal protein L8 | 6 | 3 |
| AAEL012609 | EAT35209 | $\gamma$-aminobutyric acid transaminase | 6 | 3 |
| AAEL002542 | XP_001655586 | Triosephosphate isomerase | 7 | 2 |
| AAEL006836 | XP_001652256 | Dihydropteridine reductase | 7 | 2 |
| AAEL009080 | XP_021695478 | Importin 7 | 8 | 1 |
| AAEL006670 | EAT41719 | Vitelline membrane protein 15a-3 | 8 | 0 |
| AAEL010403 | XP_001654538 | Achaete scute target 1 | 7 | 1 |
| AAEL000641 | XP_011493116 | Protein disulfide isomerase | 6 | 2 |
| AAEL022104 | ABF18250 | 60S ribosomal protein L3 | 6 | 2 |
| AAEL002861 | ABF18383 | Saccheropin dehydrogenase 1 | 7 | 1 |
| AAEL001112 | EAT47794 | Ubiquitin specific protease 5 | 8 | 0 |
| AAEL001605 | XP_001659738 | Mapmodulin/microtubule binding protein | 2 | 6 |
| AAEL013275 | XP_001663434 | Female sterile (2) ketel, importin beta | 6 | 2 |
| AAEL017315 | EJY57568 | HSC70 | 8 | 0 |
| AAEL000109 | XP_001657693 | Enolase-phosphatase E1-like | 6 | 2 |
| AAEL013857 | XP_001647837 | Serine protease immune response integrator | 1 | 6 |
| AAEL012427 | XP_001662561 | Uncharacterized protein | 7 | 0 |
| AAEL005832 | EAT42655 | Programmed cell death 4 | 6 | 1 |
| AAEL010506 | XP_001660884 | GTP-binding protein alpha subunit, gna | 1 | 6 |
| AAEL001035 | EAT47891 | Ca2+-binding protein Regucalcin/SMP30 | 6 | 1 |
| AAEL009142 | XP_001659779 | Prolyl endopeptidase | 6 | 0 |
| AAEL013284 | AAO43403 | Uncharacterized protein | 6 | 0 |
| AAEL010698 | EAT37289 | Artemis | 6 | 0 |
| AAEL001401 | XP_001659164 | Leucine-rich immune protein | 0 | 6 |
| AAEL009746 | EAT38349 | Chitinase-domain | 6 | 0 |
| AAEL007014 | EAT41361 | Odorant binding protein | 6 | 0 |
